# Catechol acetylglucose: A newly identified benzoxazinoid-regulated defensive metabolite in maize

**DOI:** 10.1101/2024.05.15.594420

**Authors:** Annett Richter, Allen F. Schroeder, Caroline Marcon, Frank Hochholdinger, Georg Jander, Boaz Negin

## Abstract

An enormous diversity of specialized metabolites is produced in the plant kingdom, with each individual plant synthesizing thousands of these compounds. Previous research showed that benzoxazinoids, the most abundant class of specialized metabolites in maize, also function as signaling molecules by regulating the production callose as a defense response. In this study, we identified catechol acetylglucose (CAG) as a benzoxazinoid-regulated metabolite that is produced from salicylic acid via catechol and catechol-glucoside. Genome wide association studies of CAG abundance identified a gene encoding a predicted acetyltransferase. Knockout of this gene resulted in maize plants that lack CAG and over-accumulate catechol glucoside. Upon tissue disruption, maize plants accumulate catechol, which inhibits *Spodoptera frugiperda* (fall armyworm) growth. Analysis of caterpillar frass showed that *S. frugiperda* detoxifies catechol by glycosylation, and the efficiency of catechol glycosylation was correlated with *S. frugiperda* growth on a catechol-containing diet. Thus, the success of *S. frugiperda* as an agricultural pest may depend partly on its ability to detoxify catechol, which is produced as a defensive metabolite by maize plants.

## Introduction

Plants produce an amazing diversity of specialized metabolites that function at all growth stages: coloring flowers and fruits to attract pollinators and frugivores ^1^, sealing the leaf surface to prevent water loss during drought ^2^, protecting the photosystem from excess light ^3^, and providing protection against herbivores and pathogens ^4^.

Benzoxazinoids are produced by grasses in the Poaceae, including maize and wheat, as well as in a few dicot genera ^5,6^. Benzoxazinoid synthesis in maize starts in the chloroplast, where 1-(2-carboxyphenylamino)-l-deoxyribulose-5-phosphate is converted to indole-3-glycerolphosphate by indole-3-glycerolphosphate synthase (IGPS) ^7^. Indole-3-glycerolphosphate is converted to indole by benzoxazinoneless 1 (BX1), and then to 2,4-dihydroxy-2H-1,4-benzoxazin-3(4H)-one (DIBOA) by four consecutive oxidation reactions catalyzed by the CYP450s BX2-BX5 ^8^. DIBOA is glycosylated by BX8 and BX9 ^9^, hydroxylated to form TRIBOA glucoside by BX6, and *O*-methylated by BX7 to form DIMBOA-glucoside ^10^. In addition to their well-established role in defense against insect herbivory ^11–13^, benzoxazinoids influence the production of other specialized metabolites, auxin metabolism, flowering time, and more ^14^.

Several plant specialized metabolites have been shown to not only have direct defensive properties but also to regulate the accumulation of other plant metabolites. Callose deposition is regulated by glucosinolate breakdown products in Arabidopsis ^15^ and benzoxazinoids in maize ^16,17^. In Arabidopsis, reduced glucosinolate biosynthesis affects phenylpropanoid production ^18^ and short-lived aglycone intermediates generated by glucosinolate hydrolysis induce jasmonate-regulated defenses ^19^. Regulatory roles were also found for volatile specialized metabolites: Arabidopsis monoterpenes induce jasmonate signaling ^20^, indole primes rice plants to herbivore attack by activating transcription of receptor-like kinases ^21^, and the green leaf volatile cis-3-hexenyl acetate primes poplar trees, elevating their defensive response to *Lymantria dispar* (spongy moth) larval feeding ^22^.

Catechol (1,2-dihydroxybenzene) is produced in bacteria ^23^ and fungi ^24^ through oxidation of salicylic acid by flavin-dependent monooxygenases ^25^. Although the presence of catechol has been reported in many plant families ^26–29^, enzymes catalyzing its biosynthesis have only recently been characterized in tomato ^30^. Plant catechol is associated with aluminum tolerance ^31,32^, affects root development when found in smoke ^33^, acts as an oviposition stimulant for *Lasioderma serricorne* (tobacco beetle) ^34^, and protects rice against *Xanthomonas oryzae* ^35^. In addition to the many functions attributed to catechol in plants, it serves as a precursor for volatile compounds, such as the flavor compound guaiacol in tomato ^28^, which can then be further modified to produce veratrole ^36^, as well as serving as a precursor for other phenolic compounds such as *O*-quinones ^37^.

In this study, we found that benzoxazinoids regulate the accumulation of catechol, catechol glucoside, and catechol acetylglucoside (CAG) in maize. Furthermore, we show that CAG is converted to catechol upon leaf damage, and that this catechol reduces *S. frugiperda* (fall armyworm) growth.

## Methods

### Plant material and growth conditions

Maize (*Zea mays* L.) seeds from the inbred line W22 and the benzoxazinoid mutant line *bx1::Ds* (AcDs-00565, B.W06.0775), were germinated on moistened filter paper in Petri dishes for three days and were then transferred into plastic pots filled with a maize soil mix [0.16 m^3^ Metro-Mix 360 (Scotts, Marysville, OH, USA); 0.45 kg finely ground lime; 0.45 kg Peters Unimix (Griffin Greenhouse Supplies, Auburn, NY, USA); 68 kg Turface MVP (Banfield-Baker Corp., Horseheads, NY, USA); 23 kg coarse quartz sand, and 0.018 m^3^ pasteurized field soil]. *Nicotiana benthamiana* plants were grown in Cornell Mix [by weight 56% peat moss, 35% vermiculite, 4% lime, 4% Osmocote slow-release fertilizer (Scotts, Marysville, OH), and 1% Unimix (Scotts, Marysville, OH)]. Maize plants used in the different assays, as well as *Nicotiana benthamiana* plants used in the overexpression assays, were grown in a temperature-controlled growth room at 23 °C, with 100 µE light intensity and a 16:8 h light:dark cycle. When grown for seeds, maize plants were grown in a temperature-controlled greenhouse ranging from 26-28 °C during the day and 24-26 °C at night, with high pressure sodium lighting supplying additional light when the photosynthetically active radiation dropped below 350 µE.

### Confirmation of a BonnMu transposon insertion

Seeds of the B73 inbred line and a BonnMu line ^38,39^ with a transposon insertion in the Zm00001d014530 gene (BonnMu0137399, from the BonnMu-7-C-0634 stock) were germinated in maize soil mix. The BonnMu line is referred to as “0634” throughout the text. Since BonnMu insertion lines harbor multiple transposon insertions, the 0634 plants were backcrossed to wildtype B73. F_2_ progeny were characterized for the transposon insertion in the gene by performing two PCR reactions (Fig. S1). The first reaction used a TIR6 oligonucleotide primer (5’ AGAGAAGCCAACGCCAWCGCCTCYATTTCGTC 3’) matching the transposon and an oligonucleotide primer complementary to the genomic sequence of the Zm00001d014530 gene (5’ AGCAACAGCAAACACTGACG 3’). The second reaction used two oligonucleotide primers deduced from the genomic sequence of maize (5’ GAGTCCCGGAGTCTGGAGAT 3’ and 5’ AGCAACAGCAAACACTGACG 3’). The PCR scheme was identical for both reactions, starting with 2 min at 95 °C, followed by 34 cycles of 30 sec 95 °C, 30 sec 58 °C, and 30 sec 72 °C. These cycles were followed by 3 min at 72 °C. A subset of plants with homozygous insertions was grown to seed-set, their progeny were grown, and siblings were cross-pollinated. The progeny of these crosses, homozygous for the transposon insertion, were used in caterpillar growth assays. The homozygous transposon-negative sibling plants went through a similar process and were used as controls when the 0634 line was used. These are referred to as B73, despite harboring an unknown number of other transposon insertions, similar to the 0634 line.

### Insect rearing conditions

*Spodoptera frugiperda* eggs were purchased from Benzon Research (Carlisle, PA, USA) and hatched on beat armyworm diet (Southland Products, Lake Village, AR, USA) in a 28 °C incubator. First-instar larvae were transferred to maize plants or artificial diet plates approximately 48 h after eggs were placed on artificial diet for hatching.

### Precursor feeding assays with catechol, resorcinol, and hydroquinone

The third leaf of W22 and *bx1* mutant plants was excised and the excision site was inserted in 2 ml tap water with a freshly prepared solution of either 200 µM catechol, 200 µM resorcinol, or 200 µM hydroquinone (Sigma Aldrich) for 24 h. Following this, the leaves were flash-frozen in liquid nitrogen and stored at -80 °C until they were used for metabolite extraction. Control leaves from *bx1* and W22 were incubated in 2 ml tap water only.

### Stable isotope labeling assays

To investigate whether indole or salicylic acid serves as a precursor for the biosynthesis of catechol derivatives, the third leaves of *bx1* and W22 plants were excised and immersed in 2 ml tap water containing either 250 µM [^13^C_8_ ^15^N] indole (Cambridge Isotope Laboratories, Andover, MA, USA) or [^13^C_6_] salicylic acid (Supelco, Bellefonte, PA) for 42 h. Subsequently, the leaves were flash frozen and stored at -80 °C until further processing for metabolite extraction.

### Genome-wide association studies

Local LD Estimation, Haplotype Inference, and Inbred Line Relationships: SNP marker data across the same GWAS diversity panel ^40^ around the most strongly associated SNP markers for each trait were downloaded from the Cyverse Discovery Environment (https://cyverse.org/, in the directory iplant/home/shared/panzea/hapmap3/hmp321) and used to estimate local linkage disequilibrium with the pairwise correlation with sliding window algorithm implemented in TASSEL 5.2.40 ^41^.

### Construct preparation for acetyltransferase overexpression

The coding region of Zm00004b012292, a predicted acetyltransferase identified by GWAS, was amplified from W22 cDNA. Specific primers were designed based on the W22_v2 maize genome: Forward primer: 5’ GGGGACAAGTTTGTACAAAAAAGCAGGCTAAaTGGCCGTGTCGCCGGACC 3’ and reverse primer: 5’ GGGGACCACTTTGTACAAAGAAAGCTGGGTTtGTCCTCTGGTGGAGCCACGC 3’. PCR amplification was performed using Advantage® Genomic LA Polymerase Mix from TaKaRa Bio USA (Mountain View, CA, USA), known for its high fidelity and GC-rich amplicons. The PCR scheme started with 1 min at 94 °C, followed by 34 cycles of 30 sec 94 °C, 30 sec 60 °C, and 1:30 min 72 °C. These cycles were followed by 5 min at 72 °C. The resulting fragment was then cloned into pDONR207 *via* the Gateway reaction (BP clonase kit, Invitrogen, Carlsbad, CA, USA) and from there into the destination vector pEAQ with the LR clonase kit (Invitrogen). Plasmids were sequenced to confirm their accuracy and were transformed to *Agrobacterium tumefaciens* strain GV3101.

### *Nicotiana benthamiana* co-infiltration with catechol

*Agrobacterium tumefaciens* strain GV3101 carrying either the pEAQ plasmid including the Zm00004b012292 gene or a *GFP* control was infiltrated into *N. benthamiana* leaves following a protocol from Sparkes *et al*. (2006) ^42^. The prepared suspensions, adjusted to an optical density at 600 nm of 0.5, were infiltrated into young, fully expanded leaves using a needleless syringe. Two days later, either 250 μM catechol or water were infiltrated into the same spots where *Agrobacterium* had been injected. Following an additional day, the tissue was harvested and immediately frozen at -80 °C for subsequent metabolite extraction.

### Artificial diet assays

Artificial diet assays were performed by placing five *S. frugiperda* neonates in plates containing control beat armyworm diet or diet supplemented with 4 µM, 10 µM, or 25 µM catechol, or 10 µM catechol glucoside. Plates were returned to a 28 °C incubator and caterpillars were allowed to feed for 5 days, after which they were weighed and survival was scored. Frass from these plates was collected for metabolite analysis, weighed and flash frozen in liquid nitrogen. Segments of the artificial diet medium were excised from the bottom of the plate, which did not come in contact with the caterpillars, and were similarly processed.

### Caterpillar feeding on maize

Caterpillar growth was assessed on the 0634 mutants and sibling B73 controls. Five *S. frugiperda* neonates were placed on two-week old plants. These were placed in micro-perforated bread bags (25.4 x 40.6 cm, PrismPak; Berwick, PA, USA) in a temperature-controlled growth room with similar conditions to those at which the maize plants were initially grown. After nine days, caterpillars were weighed and survival was scored. Caterpillar frass and caterpillars were collected from a subset of the plants, weighed and flash frozen.

### Time course of catechol accumulation and heat inactivation

Ten-day old B73 plants were used for the initial time course assay. Plant tissue (50-100 mg) was excised from young leaves and weighed. The leaves were manually ground using a pestle in 1.5 ml microcentrifuge tubes. The samples were either immediately flash frozen (0 sec) or left at room temperature for 15 sec, 45 sec, 2 min, 6 min, 18 min, or 1 h, after which they were flash frozen. When catechol was added to the samples, it was initially suspended in methanol to a concentration of 10 mM and then diluted in water to a 100 µM stock concentration. This solution was added at a 10:1 ratio to ground maize leaf samples and left for a similar duration as the non-supplemented control samples.

For a heat inactivation assay, ∼ 100 mg of the first fully expanded leaves of four-week-old 0634 and B73 plants were sampled, weighed, and ground, similar to the time course assay. A subset of samples was immediately frozen, whereas others were left for 20 min or 1 h at room temperature and were then flash frozen. Parallel samples were similarly treated, but had catechol supplemented to them as in the assay previously described. For heat inactivation following grinding, samples were sealed with Parafilm to avoid evaporation and inserted to a heating block at ∼98 °C for 30 min. Following this, samples were moved to ice for 2 min and either flash frozen or left for 20 min or 1 h at room temperature. An additional set of samples was catechol-supplemented following the 2 min on ice and left for similar durations at room temperature before flash freezing.

### Metabolite extraction and profiling

For precursor feeding assays and *N. benthamiana* overexpression assays, 100 mg samples of frozen, powdered leaf material were weighed, three volumes of 100% methanol were added to each sample, and samples were incubated with shaking for 45 min at 4 °C. Samples were centrifuged at 17,000 *g* and filtered in a 96-well MultiScreen filter plate (0.65 µm, Millipore, Burlington, MA) or centrifuged at 17,000 *g* for 10 min for a second time. For quantification an internal standard of 2-benzoxazolinone (Sigma Aldrich, St. Louis, MO) at a final concentration of 30 µM was used. For maize mutant metabolic profiling, time courses and caterpillar assays, catechol was extracted in methanol with 0.1% formic acid, to which a 2-benzoxazolineone internal standard was added to a concentration of 1.5 µg/ml. When not ground with a pestle in tubes, samples were ground using a GenoGrinder 1600 MiniG homogenizer (SPEX SamplePrep, Metuchen NJ, USA). For each mg of plant, caterpillar, or frass sample (depending on the initial weights of the sampled material, but identical for each extraction set), 4-5 µl buffer were added. Samples were vortexed and left at 4 °C for 1 h, after which they were centrifuged twice for 10 min at 14,000 *g*, and the liquid fraction was transferred to liquid chromatography glass vials.

Reverse-phase liquid chromatography was performed using a DionexUltimate 3000 Series LC system (HPG-3400 RS High-Pressure pump, TCC-3000RS column compartment, WPS-3000TRS autosampler) controlled by Chromeleon software (Thermo Fisher Scientific, Waltham, MA), and coupled to an Orbitrap Q-Exactive mass spectrometer controlled by Xcalibur software (Thermo Fisher Scientific). A Titan™ or Supelco (Sigma Aldrich) C18 UHPLC Column, 1.9 μm 100 mm × 2.1 mm was used for metabolites originating from feeding and *N. benthamiana* assays or caterpillar assays, respectively. The column temperature was set to 40 °C and the flow rate to 0.5 ml/min. Mobile phases A (H_2_O:0.1% formic acid) and B (acetonitrile:0.1% formic acid) were used for the separation of target molecular features. For precursor feeding and *N. benthamiana* assays, the gradient starting condition was 0% B at 0 min, rising to 95% B in 15 min, then was held for 0.5 min, and followed by 0.5 min re-equilibration to the starting condition. For caterpillar and time course metabolites, buffer gradients were initially set to 0% B, increasing to 15% at 7 min. Buffer B concentration was then raised to 95% at 12.5 min, held for half a minute and decreased back to 0 at 13.1 min. This concentration was held until 13.5 min. A heated electrospray ionization source (HESI-II) in negative mode was used for the ionization, with the parameters of spray voltage at 3.5 kV, the capillary temperature at 380 °C, sheath gas and auxiliary gas flow at 60 and 20 arbitrary units, respectively, S-Lens RF Level 50, and probe heater temperature 400 °C. Data were acquired in the *m/z* range of 100–900, 140,000 FWHM resolution (at *m/z* 200), AGC target 3e6, maximum injection time of 200 ms, in profile mode.

Metabolites were quantified using ThermoScientific Xcalibur Version 4.1.31.9 (Quan/Processing). For data analysis, the raw mass spectrometry files were converted to mzxml formats using the MSConvert tool ^43^ The raw mzxml files were further processed using XCMS (http://metlin.scripps.edu/download/^44^ and the CAMERA software packages for R (http://www.bioconductor.org/packages/release/bioc/html/CAMERA.html). Data were analyzed using the XCMS-CAMERA mass scan data processing pipeline ^45^. Catechol was identified by a standard (cas 120-80-9; Sigma Aldrich), as was catechol glucoside (cas 2400-71-7; BLDpharm; Cinccinati, OH, USA). CAG was identified by fragmentation patterns, feeding assays and disappearance in the 0634 mutants with parallel accumulation of catechol glucoside.

### Reversed-phase separation of methanol extracts for GC-MS

CAG was extracted from bulk snap-frozen B73 seedling leaves using 50% (v/v) methanol acidified with 0.1% (v/v) formic acid. Solid debris were removed through centrifugation and the crude extract was concentrated with a Rotovapor (Buchi). Seven ml of concentrated extracts were filled into a 25 ml round bottom flask and diluted with an additional 10 ml of methanol. Five g of silica gel (Silicycle, SiliaFlash P60 40-63um, 230-400 mesh), were added and the resulting suspension was concentrated to dryness *in vacuo*, resulting in a powder representing plant methanol extract absorbed on the silica gel. This sample was fractionated using a RediSepRf high performance C18 column (Teledyne Isco, LINCOLN, NE, USA) by reversed-phase flash chromatography using a Teledyne Isco CombiFlash RF+ Lumen instrument, with a gradient of 10-40% acetonitrile in water over 22 minutes followed by 100% acetonitrile for 11 minutes. Fractions were examined by HPLC-MS for the presence of CAG, and CAG-containing fractions were combined and concentrated to dryness in vacuo.

Following fractionation, the fractions enriched in CAG were evaporated under a nitrogen flow, resuspended in dichloromethane and derivatized using *N*-Methyl-*N*-(trimethylsilyl)trifluoroacetamide (MSTFA) containing 1% trimethylchlorosilane (TMCS; Thermo Scientific). GC-MS was performed using an Agilent (Santa Clara, CA, USA) GC-MS system (Agilent 7890B GC coupled to an Agilent 7000D triple quad mass spectrometer and a Gresel MPS2 multipurpose sampler). One µl samples were injected into an Agilent HP-5MS column (30 m x 250 µm x 0.25 µm) in spitless mode. Helium was used as the carrier gas, with a flow rate of 1.2 ml/min. The column temperature was initially set to 50 °C for 2 min and then raised to 320 °C at a rate of 20 °C/min. the column was then held at 320 °C for an additional 5 min. The mass spectra were scanned between *m/z* 40 – 900, with a scan time of 300 ms. Chromatograms were analyzed using Mass-Hunter software (https://www.agilent.com/en/promotions/masshunter-mass-spec) and compared to documented fragmentation patterns using the NIST Mass Spectral Library (https://chemdata.nist.gov/).

### Phylogenetic tree construction

For phylogenetic comparisons of maize CGA1 and similar enzymes, the following protein sequences were obtained from NCBI, either based on their homology to the maize CGA1 protein or by searching for papers which demonstrated functions for different acetyl or malonyl transferase enzymes targeting specialized metabolites as a substrate: OsMAT2 (BAD21783.1), a rice flavonoid malonyltransferase, that malonylates the 6″-hydroxyl group of the sugar ^46^; Si AOGOM (XP_004969567.1), *Setaria italica* anthocyanidin 5-O-glucoside-6’’-O-malonyltransferase; Sb A0G0M (XP_002438902.1), *Sorghum bicolor* anthocyanidin 5-O-glucoside-6’’-O-malonyltransferase; Zm PGMT1 (AQK66481.1), *Zea mays* phenolic glucoside malonyltransferase1; Nt MAT1 (NP_001312260.1), *Nicotiana tabacum* malonyl transferase, which malonylates 2-naphthol-O-glucoside on position 6 ^47^; Nto PGMT (XP_033516236.1), *Nicotiana tomentosiformis* phenolic glucoside malonyltransferase; Gh 3-O-glucoside-6-O-malonyltransferase1 (AAS77403.1), a *Glandularia x hybrida* enzyme; At5G39050 PGMT (NP_568561.4), Arabidopsis phenolic glucoside malonyltransferase, which malonylates 2-naphthol-O-glucoside on position 6 ^47^; Mt isoflavonoid malonyl transferase1 (ABY91220.1), a *Medicago truncatula* enzyme that malonylates a sugar connected to an isoflavone ^48^; At3G03480 (Z)-3-hexen-1-OL acetyltransferase (NP_186998.1), an Arabidopsis enzyme predicted by homology to acetylate alcohols ^49^; Ph benzyl alcohol benzoyltransferase (Q6E593.1), a petunia enzyme that can acetylate or add a benzoyl group to benzoyl alcohols ^50^; CrDAT deacetylvindoline 4-*O*-acetyltransferase (Q9ZTK5.1), a *Catharanthus roseus* enzyme that catalyzes the last step in vindoline biosynthesis by acetylating a hydroxyl of deacetylvindoline ^51^; Sl GAME36 (XP_004246214.1), a tomato enzyme that acetylates hydroxytomatine to form acetoxytomatine ^52^; Fa alcohol acyltransferase (AAG13130.1), a strawberry enzyme that acetylates short alcohols (C2-C7) ^53^; and Sl ASAT4 (NP_001266253.1), a tomato acyltransferase that acetylates sucrose in the biosynthesis of acylsugars ^54^.

These protein sequences were aligned using Clustal omega software (https://www.ebi.ac.uk/jdispatcher/msa/clustalo). The output of this alignment was then used to construct a maximum likelihood, midpoint-rooted tree using the IQ TREE software (http://www.iqtree.org/), with Bootstrap values that were based on 1000 replicates. The resulting tree was visualized with MEGA11 software (https://www.megasoftware.net).

### Statistical analyses

For comparisons of 0634 mutants to B73, time courses, caterpillar assays, and metabolic profiling and heat inactivation assays, JMP 16 software (SAS Institute, Cary, NC, USA) was used. For pairwise comparisons, Student’s *t*-tests were performed. For multiple comparisons a Tukey’s HSD test was used. Regression lines equality to zero was examined using PRISM 9 software (GraphPad; Boston, MA, USA) and a regression *t*-test.

## Results

Given the dual function of benzoxazinoids in direct defense and defense regulation, we initiated a search for maize metabolites that accumulate in a benzoxazinoid-dependent manner. To this end, we extracted leaf metabolites from both wildtype W22 maize and *bx1* plants, which have a mutation in the first committed step of benzoxazinoid biosynthesis. Principal component analysis showed that the overall metabolite content differs between *bx1* mutant and wildtype plants (Fig. 1A). We next searched for specific compounds that were missing in *bx1* mutants but present in the W22 wildtype plants, which would indicate regulation by benzoxazinoids and possible functions in plant protection from insect herbivores. In an analysis of the chromatographic peaks, we found two metabolites with a common *m/z* 109.02 breakdown product. The first peak had a 271.08 *m/z*, indicating a possible addition of a glucose to the *m/z* 109.02 mass feature (Fig. 1B), whereas the second peak had an *m/z* of 313.09, indicating a possible acetylation of the first compound (Fig. 1C). The exact mass of the *m/z* 109.02 feature suggested the presence of a benzene ring with two hydroxylations. There are three compounds that differ in the hydroxylation sites on the benzene ring but have the same mass: catechol (1,2-benzenediol), resorcinol (1,3-benzenediol), and hydroquinone (1,4-benzenediol). To determine whether one of these compounds was a precursor of the two metabolites that were co-synthesized with benzoxazinoids, we fed them to the *bx1* mutants and examined whether the *m/z* 271.08 and *m/z* 313.09 peaks, which are not normally present in the *bx1* mutants, would appear. Catechol feeding restored these two peaks (Fig. 2), whereas feeding of both hydroquinone or resorcinol led to the formation of *m/z* 271.08 and *m/z* 313.09 peaks that were not present in *bx1* and had different retention times compared to the native W22 mass features. We examined the retention time and fragmentation pattern of a catechol glucoside standard and found that it was identical to that of the compound with *m/z* 271.08 that we had identified. This suggested that the *m/z* 313.09 peak was an acetylated version of catechol glucoside.

**Figure 1.**
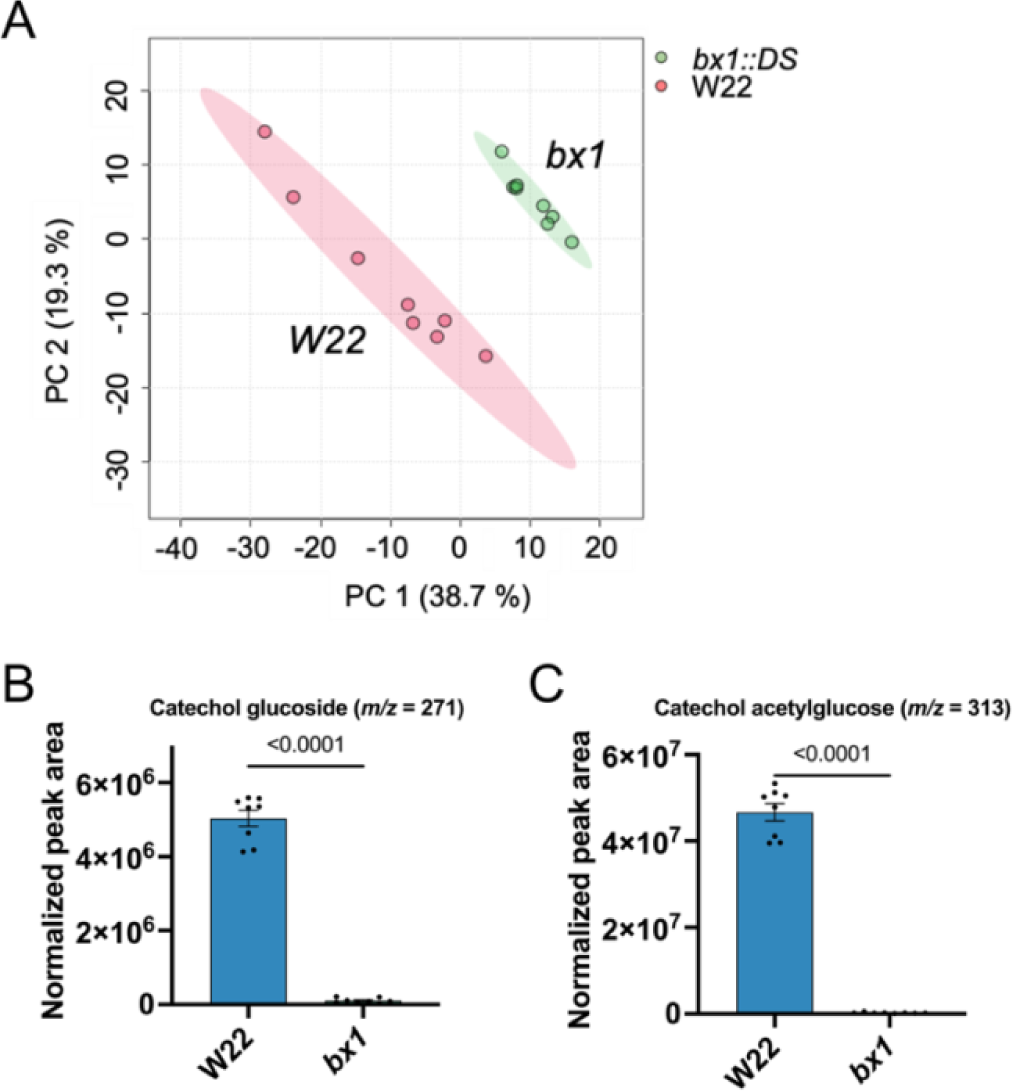
Metabolic comparison of W22 wildtype and *bx1* mutant maize. (A) Principal component analysis of untargeted metabolite profiling data. Raw metabolite data from HPLC-MS in negative ionization mode are in Table S1. The graph was made with Metaboanalyst. (B) Normalized peak area of catechol glucoside (*m/z*= 271.08) in wildtype and *bx1* mutant plants. (C) Normalized peak area of catechol acetylglucose (*m/z*= 313.09) in wildtype and *bx1* mutant plants. P values were determined by two-tailed Student’s *t*-tests. Mean +/− standard error (s.e). of N = 8.

**Figure 2.**
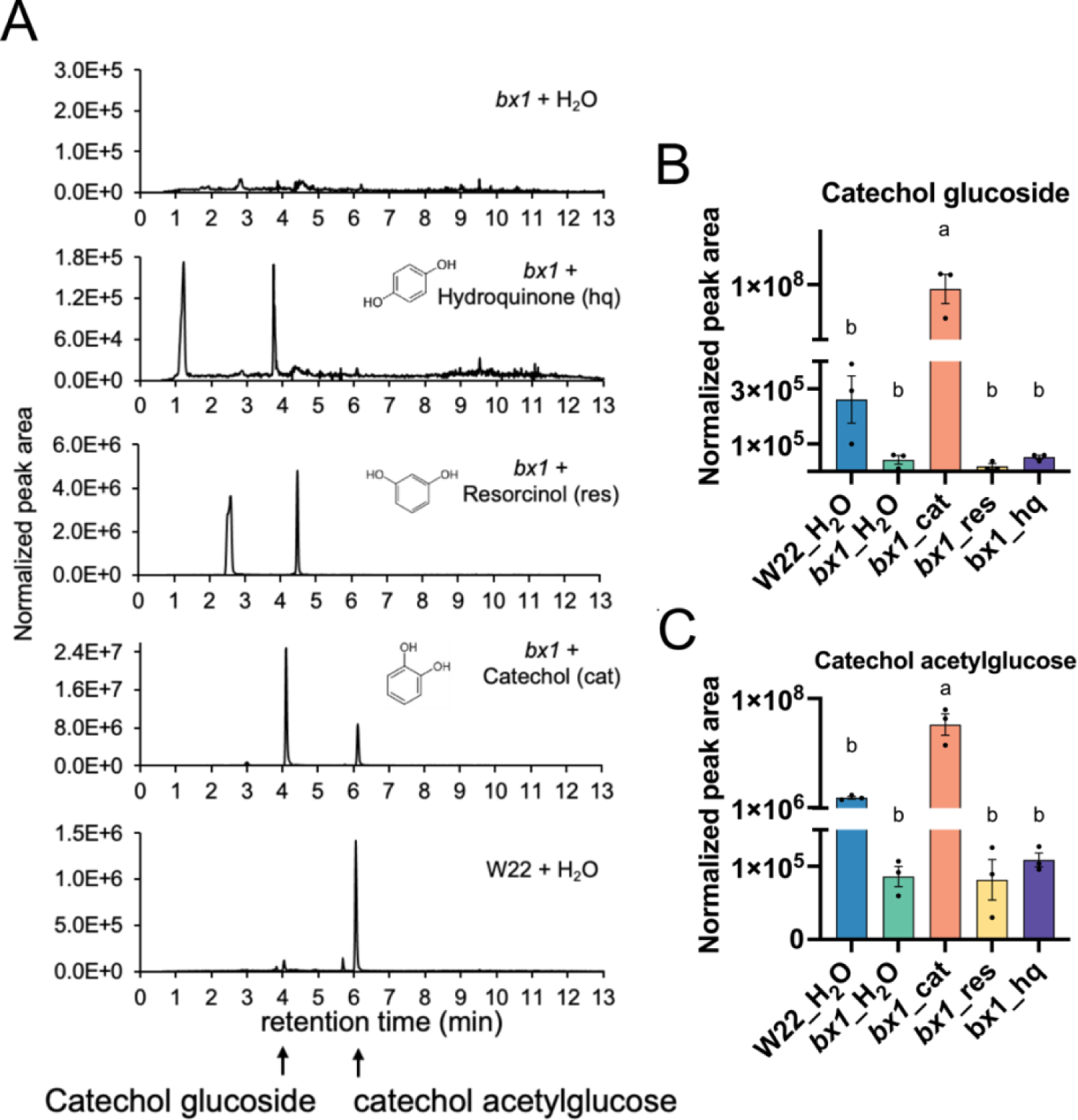
Hydroquinone, resorcinol, and catechol feeding experiment with bx1 mutants. (A) Chromatograms from the feeding experiment, showing accumulated compounds with *m/z* 109.02, corresponding to fragments of catechol glucoside and catechol acetylglucose. (B) catechol glucoside and (C) catechol acetylglucose, normalized peak area, comparing treatment with H_2_0 (control), hydroquinone (hq), resorcinol (res), and catechol (cat). Wildtype W22 is also shown as a control. Different letters indicate a significance of p < 0.05 as determined in a Tukey’s HSD test. Mean +/− s.e. of N = 3.

To determine whether catechol glucoside is produced from indole, the product of the BX1 enzyme, we fed [^13^C_8_ ^15^N] indole to *bx1* mutant leaves. This increased the abundance of unlabeled catechol glucoside (Fig. 3A) and CAG (Fig. 3C), but no labeled catechol glucoside with an expected *m/z* 277.1 (Fig. 3B) or CAG with an expected *m/z* of 319.11 (Fig. 3D) were formed. By contrast, feeding [^13^C_8_ ^15^N] indole to *bx1* mutants increased the abundance of isotope-labeled DIMBOA glucoside (Fig. 3E), but not that of unlabeled DIMBOA glucoside (Fig. 3F). These results indicate that indole is incorporated directly into DIMBOA, which then regulates the synthesis of catechol glucoside. However, indole, benzoxazinoids, or their breakdown products are not directly incorporated into catechol glucoside.

**Figure 3.**
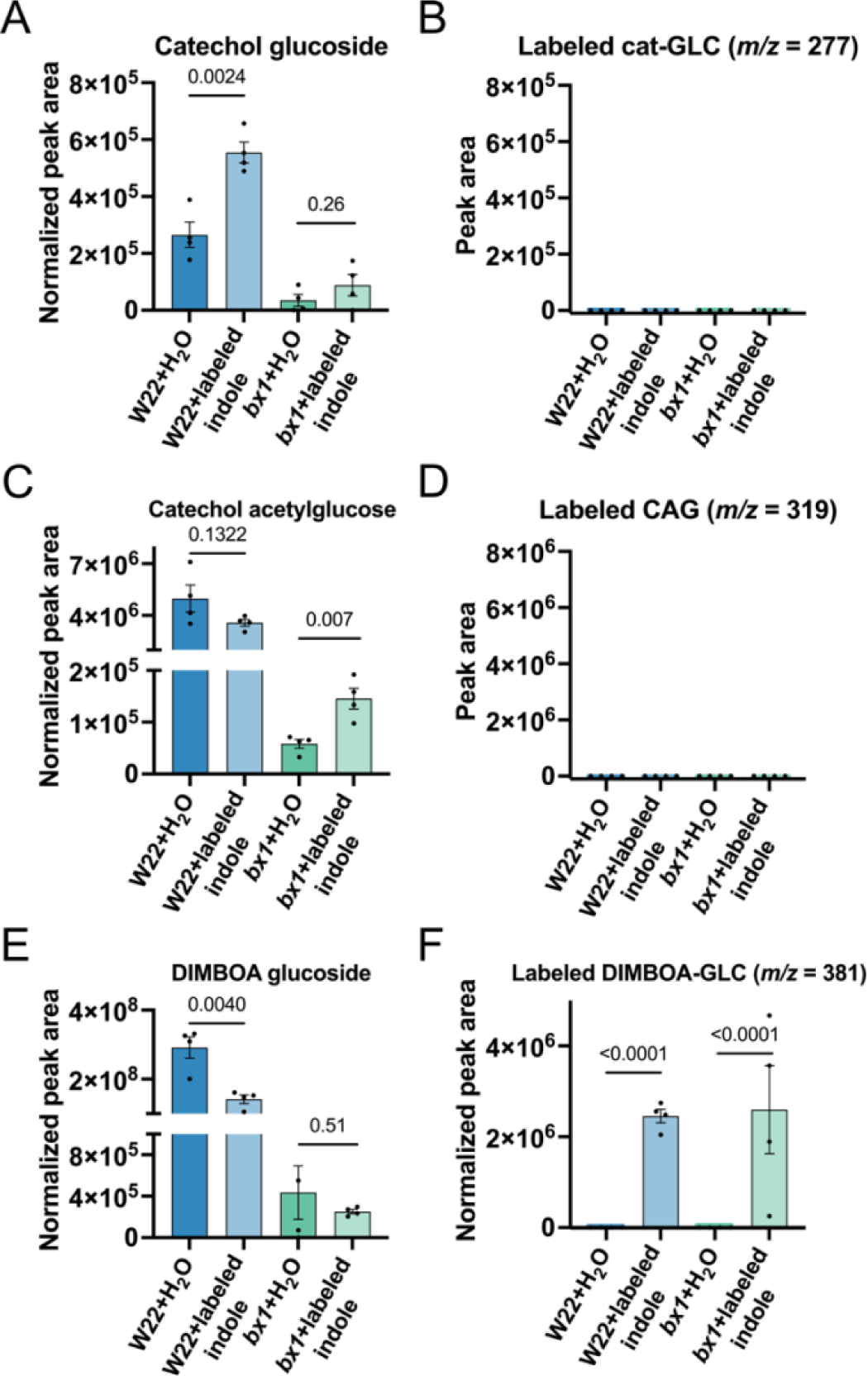
Benzoxazinoid and catechol derivatives after [^13^C_8_ ^15^N] indole feeding of W22 wildtype and *bx1* mutant maize. Normalized peak areas of (A) unlabeled catechol glucoside (*m/z* 271.08), (B) labeled catechol glucoside (predicted *m/z* 277.1, (C) unlabeled catechol acetylglucose (*m/z* 313.09), (D) labeled catechol acetylglucose (predicted *m/z* 319.11), (E) unlabeled DIMBOA-glucoside (*m/z* 372.09), and (F) labeled DIMBOA-glucoside (*m/z* 381.11). Mean +/− s.e. of N = 4. P values were determined by comparing labeled and control samples using two-tailed Student’s *t*-tests.

Salicylic acid is a precursor for catechol synthesis in bacteria, fungi, and tomatoes ^23,24,30^. Feeding [^13^C_6_] salicylic acid led to formation of labeled catechol glucoside (Fig. 4B) and the acetylated compound (Fig. 4D) in both W22 plants and *bx1* mutants. By contrast, there was no increase in the corresponding unlabeled compounds (Fig. 4A, C), indicating that salicylic acid is directly incorporated, rather than regulating their biosynthesis.

**Figure 4.**
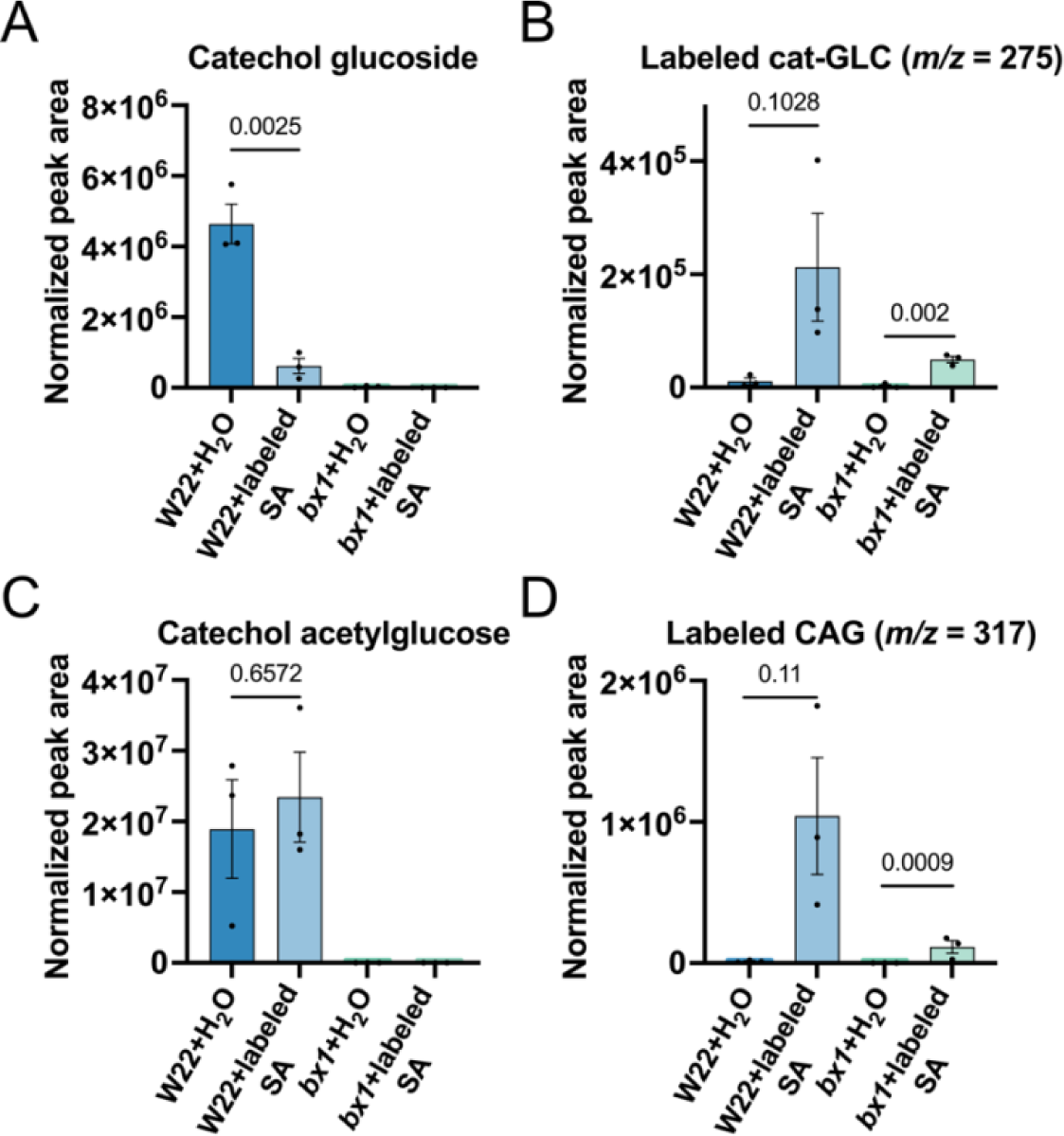
Labeling of catechol derivatives with deuterated salicylic acid. Normalized peak areas of (A) unlabeled catechol glucoside (*m/z* 271.08), (B) labeled catechol glucoside (*m/z* 275.1), (C) unlabeled catechol acetylglucose (m/z 313.09), (D) labeled catechol acetylglucose (CAG) (*m/z* 317.11). Mean +/− s.e. of N = 3. P values were determined by comparing labeled and control samples using two-tailed Student’s *t*-tests.

To identify genes regulating the biosynthesis of the *m/z* 271.08 and *m/z* 313.09 compounds, we conducted genome-wide association studies (GWAS) using a previously published metabolomics dataset ^40^. We found a locus on chromosome 5 that was significantly correlated with abundance of the *m/z* 313.09 compound, and contained a gene (Zm00001d014530) predicted to have an acetyltransferase function (Fig. 5A). To obtain insight into whether the Zm00001d014530 gene was involved in maize responses to insect herbivory, we examined its transcriptional response to jasmonate treatment and caterpillar feeding, using published datasets of RNA-seq following these treatments ^55,56^. Jasmonate treatment led to a substantial increase in Zm00001d014530 transcript abundance (Fig. 5B). Feeding of second to third instar *Spodoptera exigua* caterpillars confined to maize leaves in clip cages led to an increase in the transcript abundance starting at the first time point measured, at 1 h post feeding, and up the last measurement at 24 h post feeding (Fig. 5C).

**Figure 5.**
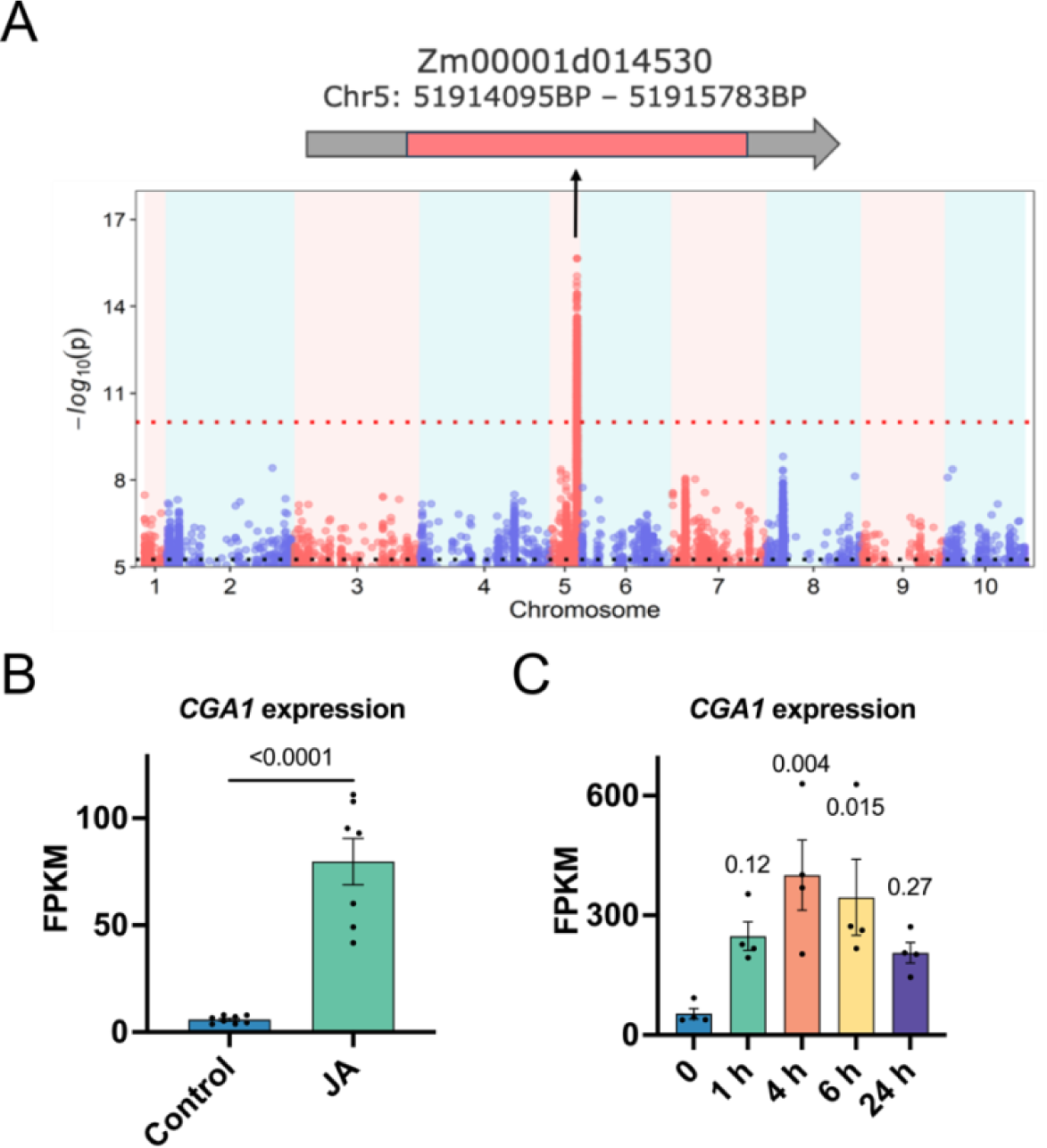
Identification and induced expression of a catechol glucoside acetyltransferase. (A) Genome wide association study (GWAS) of catechol glucoside (*m/z* 271.08) abundance. Red dotted line highlights the highest peak. Black dotted line indicates genome-wide significance threshold line. (B) Induction of candidate acetyltransferase expression by jasmonic acid treatment of wildtype W22 (data from Feiz *et al*, 2024 ^56^), mean +/− s.e. of N = 8. (C) Induction by *Spodoptera exigua* feeding over 24 hours in inbred line B73 (data from Tzin *et al*, 2017 ^55^). Mean +/− s.e. of N = 4. P values were determined using a two-tailed Student’s *t*-test (B) or Dunnett’s test (C) compared to the B73 control.

To examine whether the Zm00001d014530 gene was involved in the synthesis of the *m/z* 313.09 compound, we transiently expressed it in *Nicotiana benthamiana* and obtained a maize transposon insertion line with a knockout of this gene. Feeding *N. benthamiana* plants with catechol alone led to formation of catechol glucoside (Fig. 6A, B) but not the *m/z* 313.09 compound (Fig. 6A, C). By contrast, feeding with both catechol and transiently expressing the predicted acetyltransferase gene reduced the abundance of catechol glucoside (Fig. 6A, B) and led to accumulation of the *m/z* 313.09 compound (Fig. 6A, C). Following validation of homozygous transposon insertions in the mutant line (Fig. S1), we found that catechol glucoside abundance was increased (Fig 6E) and abundance of the *m/z* 313.09 compound was reduced to near zero levels (Fig. 6F), indicating that Zm00001d014530 alone is responsible for the acetylation of catechol glucoside and that the catechol glucoside was the direct precursor of the *m/z* 313.09 compound. Given the encoded enzymatic function, we have named the Zm00001d014530 gene *catechol glucoside acetyltransferase 1* (*Cga1*).

**Figure 6.**
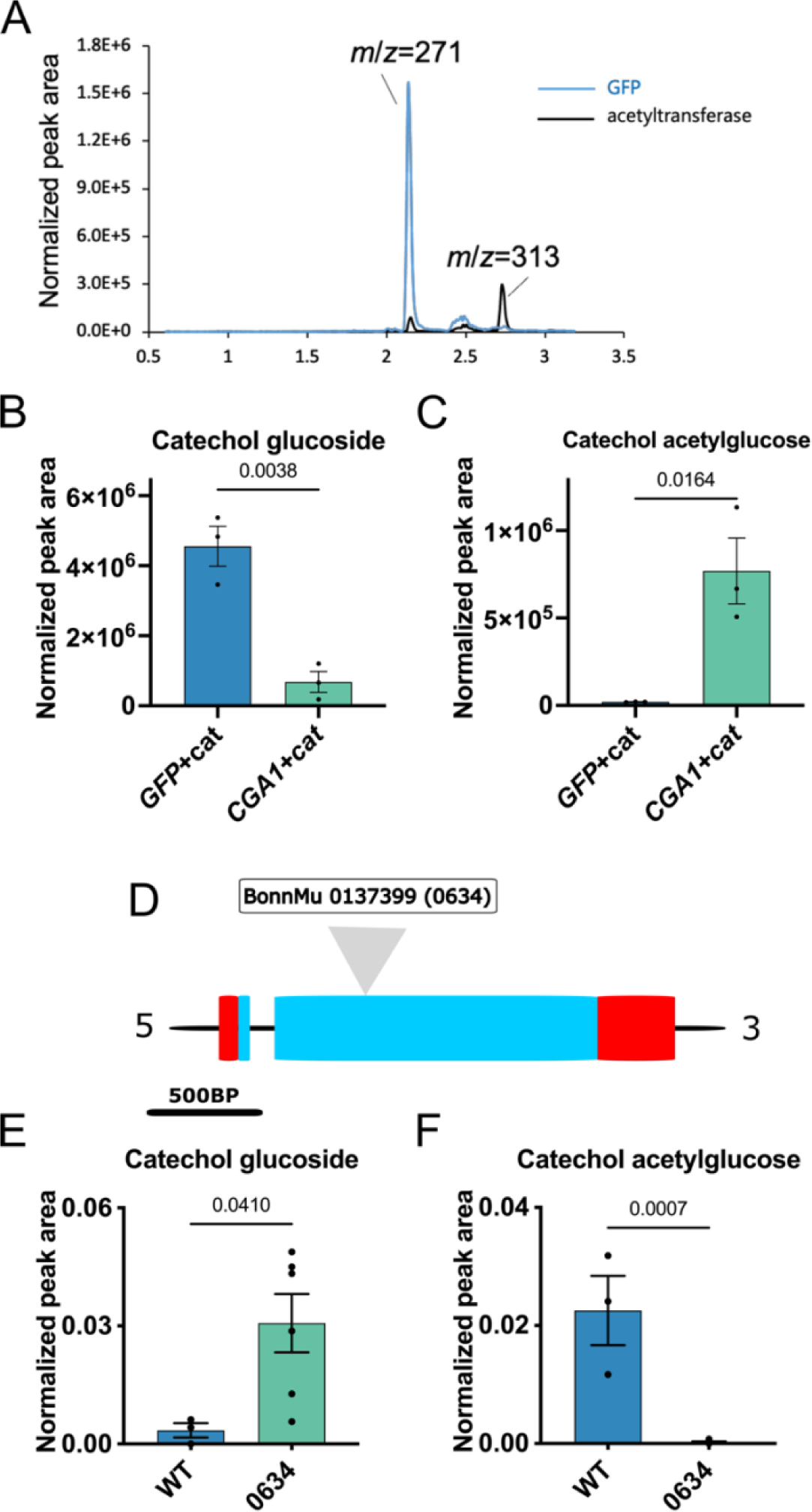
Characterization of a catechol glucoside acetyltransferase. (A) HPLC-MS chromatogram showing transient expression of a maize acetyltransferase in *Nicotiana benthamiana*, with catechol co-infiltration. A *GFP*-expressing construct was used as the control. (B) and (C) show quantification of catechol glucoside and catechol acetylglucose, respectively. Mean +/− s.e. of N = 3. (D) The structure of Zm00001d014530 (the B73 *CGA1* gene) and the location of the transposon insertion in the 0634 mutation. Red boxes indicate untranslated regions and the blue box indicates an exon. Abundance of catechol glucoside (E) and catechol acetylglucose (F) in wildtype-B73 and 0634 mutant plants, as determined by LC-MS analysis. P values were determined by a Student’s *t*-test. (E-F) WT n=3, 0634 n=6.

To assess whether the acetylation occurs on the glucose or on the second hydroxyl of the catechol, we performed a phylogenetic analysis of acetyltransferase proteins that were demonstrated to acetylate specialized metabolites, or proteins homologous to them. Zm00001d014530 clustered with enzymes that acetylate or malonylate sugars connected to phenolic compounds (Fig. S2). Of these, the enzymes from rice, Arabidopsis, tobacco, and *Medicago truncatula* had experimentally verified functions. Enzymes acetylating sucrose, alcohols, non-glycosylated phenolics, precursors of vinblastine, and hydroxytomatine did not cluster with this group.

To further verify the site of acetylation, we analyzed the fragmentation of a fraction enriched with the *m/z* 313.09 compound by GC-MS. Due to lack of both a standard and library entries, we cannot be certain about the fragmentation of the *m/z* 313.09. However, the enriched fragment had the highest fragmentation similarity (R match) to a methylated acetylglucose (Fig. S3). This fragment’s most prominent mass feature had an *m/z* of 204, similar to the mass of an acetylated glucose which had lost a hydroxyl where it was connected to the catechol. An additional prominent peak had an *m/z* of 217, as expected for methylated acetylglucose that had lost a hydroxyl. It is possible that this methyl group is part of the catechol, which was connected to acetylglucose and broke upon ionization.

Since catechol glucoside production was regulated by benzoxazinoids (Fig. 1), and *Cga1* transcription was induced by both jasmonate and caterpillar feeding (Fig. 5B,C), it seemed likely that catechol has a defensive function in maize. We performed artificial diet experiments with both catechol and catechol glucoside, though not with CAG, which is not commercially available. This experiment was based on the assumption that, similar to many defensive specialized metabolites ^52,57^, glycosylation and acetylation are likely to be mechanisms to store catechol as a non-toxic precursor. Both maize and *N. benthamiana* rapidly form catechol glucoside after catechol infiltration. To perform an artificial diet experiment using biologically relevant concentrations, we generated catechol and catechol-glycoside calibration curves and determined that W22 wildtype maize leaves contained approximately 4 µM catechol-glycoside and CAG. Since catechol is volatile, we assumed that its concentration in plates would decrease over time, and therefore added, in addition to 4 µM, both 10 µM and 25 µM catechol, as well as 10 µM catechol-glycoside which we found to be more stable.

*Spodoptera frugiperda* feeding on plates containing 25 µM catechol displayed significantly reduced growth (Fig. 7A), although survival remained unaffected (Fig. 7B). In the frass of caterpillars from the different treatments, an *m/z* 109.02 peak was similar across all treatments, indicating an endogenous compound produced by the caterpillars (Fig. 7C). By contrast, the abundance of catechol glucoside in the frass increased as the catechol concentration in the plates increased, indicating that the caterpillars were metabolizing catechol to catechol glucoside (Fig. 7D). We examined the correlation between catechol, catechol glucoside, their ratio, and the caterpillar weight of large and small caterpillars fed on 25 µM catechol. This showed significant negative correlations among two of the three factors (Fig. 7E-G), indicating that caterpillars that were able to metabolize catechol to catechol glucoside more efficiently grew to a larger size.

**Figure 7.**
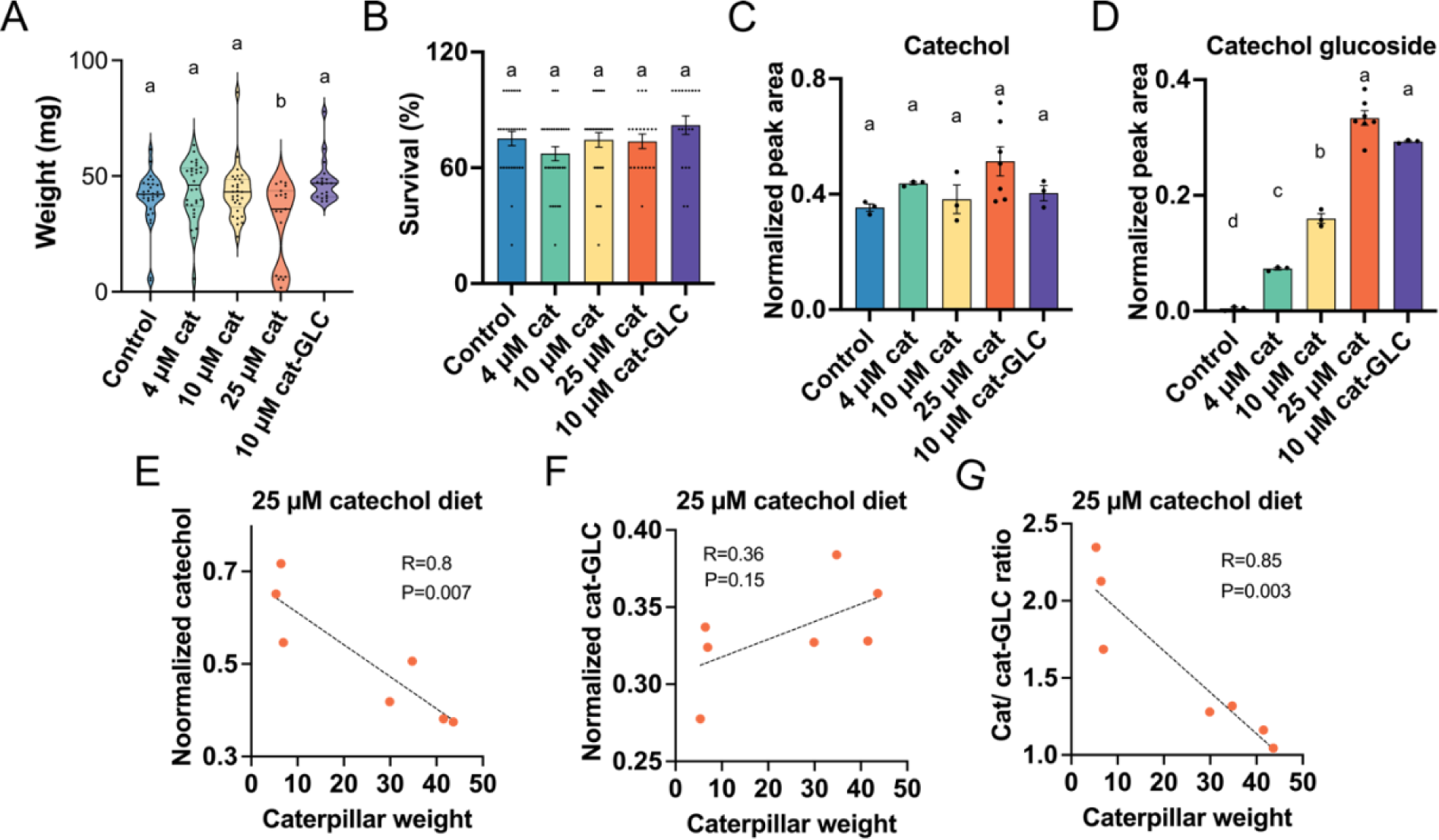
Effects of catechol feeding on Spodoptera frugiperda growth, and catechol detoxification. (A-B) *S. frugiperda* weight (A) and survival (B) following feeding on artificial diet supplemented with rising catechol concentrations, or catechol glucoside. (C-D) Metabolite abundance in *S. frugiperda* frass following feeding on the artificial diet. (C) Catechol and catechol-like compounds co-eluting in the frass chromatogram. (D) Catechol glucoside. (E) correlation of caterpillar weight to catechol and co-eluting catechol-like compounds. (F) Correlation of caterpillar weight with catechol glucoside. (G) Correlation of caterpillar weight with the ratio of catechol and catechol glucoside. (A-B) N = 19-30. (C-D) N = 3-8. (E-G) N = 8. Different letters indicate a significance of p < 0.05, as determined by a Tukey’s HSD test. P values which appear on scatter plots indicate significance of slopes equality to zero, as determined with a regression *t*-test. Cat – catechol, cat-GLC – catechol glucoside.

Given these findings, we aimed to determine whether catechol glucoside and/or CAG were being converted to catechol upon caterpillar feeding, and how fast catechol would be metabolized back to catechol glucoside. We ground 10-day old B73 maize seedling at 23 °C and incubated the samples at 23 °C for increasing periods of time up to 1 hour, with and without added catechol, before stopping enzymatic reactions by flash freezing the samples in liquid nitrogen. Examining abundance of catechol, catechol glycoside, and CAG in the samples revealed that following grinding, free catechol was formed after one hour of incubation. In the catechol-supplemented samples, we found that as time passed the concentration of catechol was reduced, until the beginning of endogenous catechol formation at 1 h after grinding (Fig. 8D). These contrasting trends of catechol formation and disappearance with passing time could also be seen in significant correlations between time after grinding and catechol abundance in caterpillar-fed and unfed samples (Fig. 8G-H).

**Figure 8.**
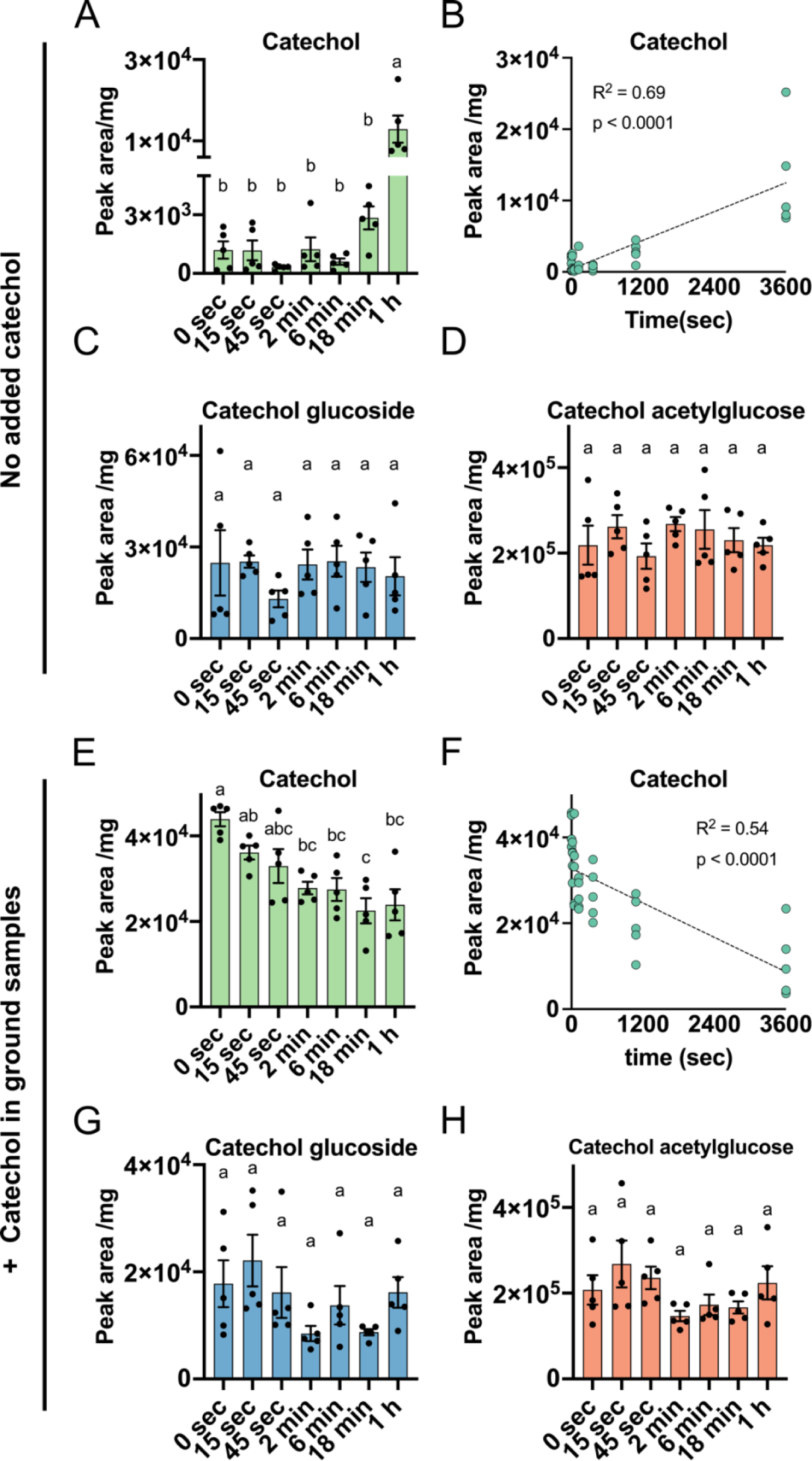
Effects of incubation duration on catechol and catechol derivative content of ground B73 samples, and samples to which 10 µM catechol were added. (A-D) No supplemented catechol. (E-H) following catechol supplementation. (A, E) Catechol abundance. (C, G) Catechol glucoside abundance. (D, H) CAG abundance. (B, F) Correlation of catechol content to incubation time. (B) Un-supplemented samples. (F) Catechol supplemented samples. N = 5. Different letters indicate a significance of p < 0.05 as determined by a Tukey’s HSD test. P values that appear on scatter plats indicate significance of the slope’s equality to zero as determined by a regression *t*-test.

As catechol in liquid solution may be oxidized in a non-enzymatic manner, we asked whether this reduction in catechol abundance with time was facilitated by an enzymatic process. We ground maize tissue, heated it to inactivate enzymes and incubated the samples with and without supplemented catechol. Whereas catechol in non-heat-treated samples became less abundant over time (Fig. S4A,G), this did not happen in heat-treated samples (Fig. S4D, H), Furthermore, the ratio between catechol, catechol glucoside, and CAG changed drastically upon heating. Prior to heating (without incubation), only traces of catechol, small amounts of catechol glycoside, and substantial amounts of CAG could be detected in maize leaves. This can be seen in the pre-heated ratios, which lean heavily towards CAG and catechol glycoside (Fig. S4I-K). However, following heating these ratios are reduced, indicating that heating leads to conversion of CAG to catechol glycoside and catechol.

Since *S. frugiperda* growth was reduced by catechol in artificial diet assays, we determined whether caterpillars would grow better on the 0634 acetyltransferase mutants, which have nearly no CAG, but increased catechol glucoside abundance. We found that *S. frugiperda* grew and survived similarly on the mutant and wildtype plants (Fig. 9A,D). When examining the abundance of catechol glucoside and CAG in the frass and the bodies of caterpillars from wildtype and mutant maize plants, as well as those raised on artificial diet, we found that, similar to the frass from artificial diet assays, both caterpillars and their frass contained catechol glucoside. Interestingly, CAG was present at high levels in the frass of caterpillars from wildtype maize, indicating that this compound is not completely metabolized in the caterpillar gut. In addition, we examined catechol, catechol glucoside, and CAG abundance in wildtype and mutants, ground and incubated for different periods, with and without catechol supplementation, to determine whether the acetyltransferase mutation would affect the rate of catechol formation and removal. Here we found that mutants and wildtype displayed similar dynamics (Fig. 9G-I).

**Figure 9.**
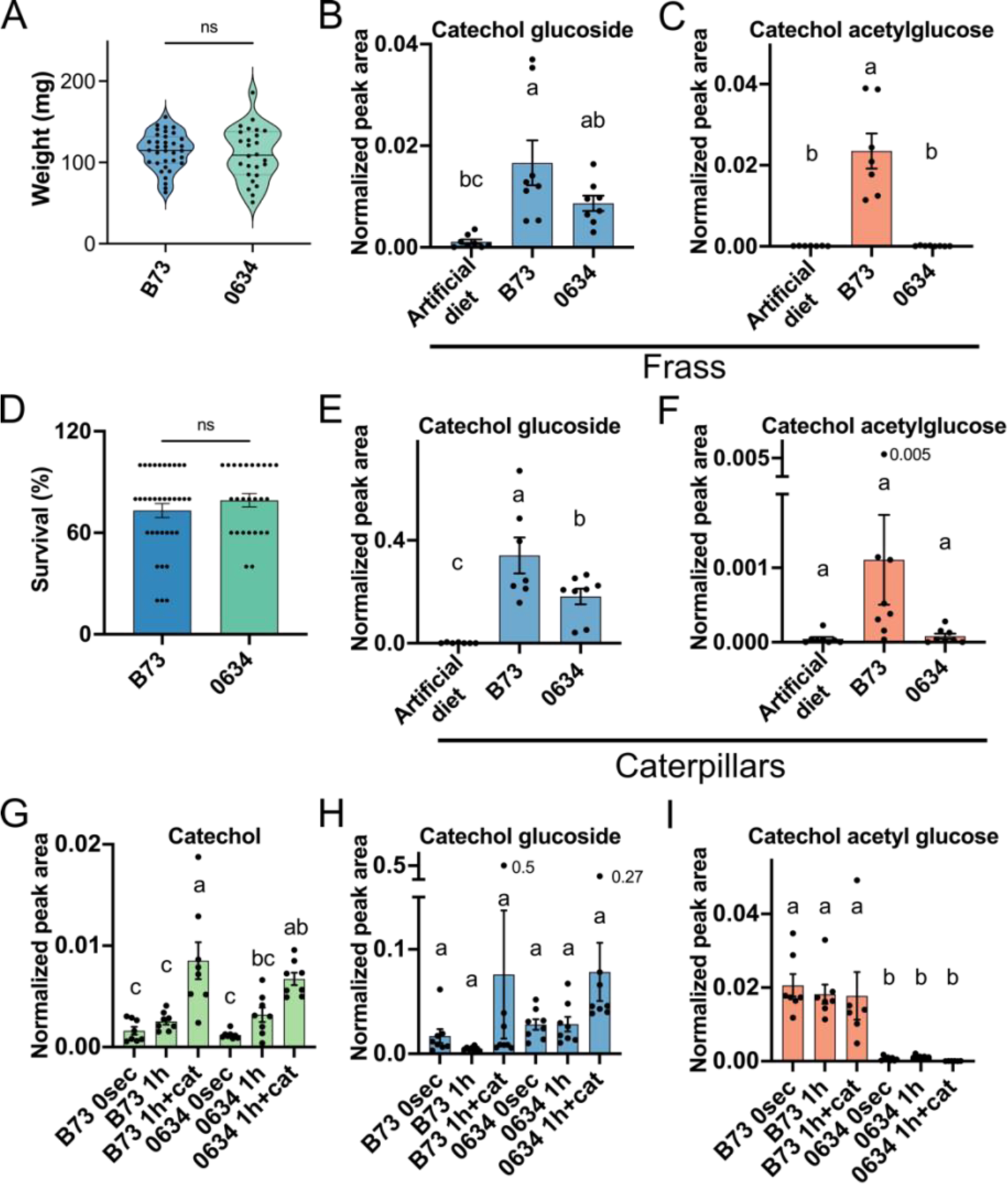
Effects of maize catechol acetylglucose production on Spodoptera *frugiperda* growth, survival and metabolite detoxification. (A) Caterpillar weight and survival (D) when feeding on B73 wildtype plants or 0634 mutants which do not produce CAG. (B-C) Catechol glucoside (B) and CAG (C) abundance in *S. frugiperda* frass following feeding on an artificial diet not containing catechol, on B73 plants or on 0634 mutants. (E-F) Catechol glucoside and CAG abundance of *S. frugiperda* caterpillars following feeding on an artificial diet not containing catechol, on B73 plants, or on 0634 mutants. (G-I) Metabolite abundance of ground B73 and 0634 leaves not incubated (0 sec) incubated for 1 h after grinding and supplemented with catechol following grinding and incubated for 1 h. (G) catechol. (H) Catechol glucoside. (I) CAG. (A,D) N = 26-35. (B-C), N = 7-8. (E-F) N = 8. (G-I) N = 6-8. “ns” indicated no significant difference between samples as determined by a Student’s *t*-test. Different letters indicate a significance of p < 0.05 as determined by a Tukey’s HSD test.

## Discussion

Plants synthesize an enormous variety of specialized metabolites. However, in many cases, it remains unknown which ones function in interactions with the environment, are biosynthetic intermediates, or are inactivated and stored for future use. We have identified two compounds, catechol glucoside and catechol acetylglucose, whose biosynthesis is regulated by the presence of benzoxazinoids. Whereas catechol inhibits caterpillar growth, catechol glucoside does not. Maize plants form catechol upon mechanical damage, but at the same time re-glycosylate the released catechol to its inactive form. In addition, we found that *S. frugiperda* growth is inhibited by catechol, and that this insect metabolizes catechol to catechol glucoside, while likely avoiding absorbance of catechol acetylglucose, which is secreted in the frass.

For many years, despite knowledge of catechol’s presence in plants ^26–29^ and its synthesis mechanisms in bacteria ^23^, genes converting salicylic acid to catechol in plants were not identified. This changed when Zhou *et al*. (2021) demonstrated that salicylic acid 1-hydroxylase forms catechol from salicylic acid in tomato ^30^. This characterization of a hydroxylase responsible for catechol synthesis led us to hypothesize that catechol synthesis in maize would take a similar path, and demonstrated this by feeding *bx1* mutants with stable isotope-labeled salicylic acid. While our GWAS analysis successfully identified the acetyltransferase responsible for catechol acetylglucose production, no other genes were correlated with this metabolite (Fig. 5A). Furthermore, despite not identifying the immediate target of benzoxazinoid regulation of catechol synthesis, we used indole labeling experiments to show that this regulation is not through the direct breakdown of benzoxazinoids to catechol.

One major question we dealt with throughout this study was: What is the function of CAG itself? Is it toxic to herbivores or is the acetylation part of its detoxification process? Some specialized metabolites are modified to reduce their toxicity to plants prior to herbivore feeding. For instance, benzoxazinoids are glycosylated to reduce their autotoxicity ^58^. In other cases, glycosylation is necessary to enable transport to the vacuole for storage or degradation ^57^. Interestingly, as part of the detoxification pathway of α-tomatine, this metabolite is first acetylated and only then glycosylated to form the less toxic esculeoside A ^52^.

Plants use many different methods to avoid harming themselves while harming herbivores that feed upon them. One mechanism is targeting metabolic systems that exist only in the animals. Nicotine targets acetylcholine receptors in the nervous system ^59^, capsaicin targets mammalian vanilloid receptors ^60^, and cardenolides target animal sodium-potassium ATPases ^61^. All these would not need to be detoxified as they do not harm the plant. By contrast, cannabinoids, despite targeting nervous system receptors, are compartmentalized in glandular trichomes. The reason for this localization could be autotoxicity of precursors such as daurichromenic acid and grifolic acid ^62^. This emphasizes the complex regulation plants employ to produce, store and utilize what are essentially poisons without harming themselves. It is difficult to determine from our results whether the acetylation of catechol glucoside is a method of detoxification or has other functions in regulating catechol and its derivatives. On one hand, *S. frugiperda* was unaffected by the complete elimination of CAG and parallel accumulation of catechol glucoside in the maize knockout line. On the other hand, *S. frugiperda* pass catechol acetylglucose intact through their digestive system, indicating that this may be a coping mechanism for a compound that would be toxic to different insects not adapted to feeding on maize.

In the never-ending evolutionary arms race of plants and their herbivores, plants accumulate defensive compounds, while their herbivores acquire mechanisms to continue consuming the plants. Examples of this struggle can be seen in plants that evolved several potent defensive compound classes. *Erysimum cheiranthoides* accumulates both glucosinolates and cardenolides ^63^. When cardenolide biosynthesis is eliminated, some *Brassicaceae-*specialist and generalist herbivores perform better on the plants, whereas others are unaffected, demonstrating the advantage of acquiring this additional defensive layer as well as the continuous necessity to develop novel defenses ^63^. Similarly, *N. benthamiana* produces both nicotine and acylsugars. When these two compounds are knocked out individually and in combination, different herbivores are affected. However, there remain herbivores that are tolerant to both of these defenses ^64^.

*Spodoptera frugiperda* is naturally a herbivore of maize plants ^65^. Despite its origin in the Americas, it has recently spread as an introduced species to the much of the rest of the world, including Africa, Asia, and Australia ^66,67^, where it causes substantial damage to maize crops ^68–70^. Despite feeding on a broad range of plant species, *S. frugiperda* performs better on maize compared to sorghum, rice, and wheat ^71^ and prefers to oviposit on maize ^72^. Furthermore, *S. frugiperda* was found to suppress maize induction of herbivore-induced plant volatiles in maize and to do this to a greater extent than other generalist caterpillars ^73^. When comparing the damage *S. frugiperda* causes to crops worldwide, its effects on maize were the most widespread and led to some of the most substantial crop losses ^74^.

When analyzing genes that may contribute to metabolite detoxification by *S. frugiperda*, glycosyltransferases were found to be overrepresented in this species ^75^. More specifically, it was shown that benzoxazinoids, similar to catechol, are stored in their conjugated inactive form in the plant and are activated upon insect feeding. These compounds were found to be glycosylated in the *S. frugiperda* gut and were subsequently detected in their glycosylated form in frass ^76^. Glycosylation as a form of detoxification of catechol, has been demonstrated in other caterpillar species. *Lymantria dispar* feeding on catechol-rich poplar plants detoxified catechol by three mechanisms, including two forms of glycosylation ^77^. In contrast to this detoxification method, Schnurrer et al. (2023) ^78^ found a different approach in detoxification of salicortinoids, including catechol, in six specialist caterpillars. In this case, salicortin breaks down to salicin and catechol at a pH similar to that in the caterpillar gut. However, the specialist caterpillars reduce salicortin to salicortinol, which does not form catechol ^78^. It is interesting to note that *S. frugiperda*, which is a broad generalist, as well as being a major maize pest, uses a mechanism seen in generalist rather than specialist caterpillars.

In conclusion, CAG is a novel maize metabolite whose biosynthesis is benzoxazinoid-regulated. A previously uncharacterized acetyltransferase converts catechol-glucoside to CAG, and mutants in this gene do not produce CAG while accumulating more catechol glucoside. CAG breaks down to catechol following tissue disruption, and this catechol impedes *S. frugiperda* growth. However, *S. frugiperda* is able to detoxify catechol by glycosylation, and at biological concentrations the caterpillars grow normally. Elevating catechol concentrations revealed that caterpillars that were not able to detoxify catechol to catechol glucoside, grew at slower rates, explaining both why maize plants accumulate CAG and providing a further explanation for *S. frugiperda*’s success in feeding on maize plants.

## Acknowledgements

The research was funded by United States Department of Agriculture – National Institute of Food and Agriculture awards 2022-67012-36739 to BN and 2021-67014-342357 to GJ, subcontracts from US National Science Foundation awards 2019516 and 2019674 to GJ, and Deutsche Forschungsgemeinschaft (DFG) award MA8427/1-1 to CM.

## Author contributions

AR designed and performed experiments and analyzed the data. AS performed experiments. CM and FH developed and supplied the BonnMu transposon insertion line. GJ conceived of the project, supervised the research, and revised the manuscript. BN designed and performed experiments, analyzed data, and wrote the manuscript. All authors revised and approved the final draft of the manuscript.

## List of supplemental materials

**Figure S1.** PCR amplification of the *Cga1* gene and a combination of the *Cga1* gene and the transposon *TIR6* sequence from a B73 x 0634 maize F_2_ progeny.

**Figure S2.** Phylogenetic tree of acetyltransferases and malonyltransferases with different substrates.

**Figure S3.** Gas chromatography-mass spectrometry analysis of the *m/z* 313.09 compound fragmentation.

**Figure S4.** Effects of incubation time and heat inactivation on the abundance of catechol metabolites, with and without catechol supplementation.

**Table S1.** Raw data for Figure 1A

**Table S2.** Raw data for Figure 1B,C

**Table S3.** Raw data for Figure 2

**Table S4.** Raw data for Figure 3

**Table S5.** Raw data for Figure 4

**Table S6.** Raw data for Figure 5A

**Table S7.** Raw data for Figure 5B,C

**Table S8.** Raw data for Figure 6E,F

**Table S9.** Raw data for Figure 7

**Table S10.** Raw data for Figure 8

**Table S11.** Raw data for Figure 9

**Table S12.** Raw data for Figure S4

**Supplemental Data S1**. FASTA file of protein sequences used to generate the phylogenetic tree of acetyl and malonyl transferase enzymes.

