## Supplemental Figures S1-S4 for "Catechol acetylglucose: A newly identified benzoxazinoid-regulated defensive metabolite in maize"

**Supplementary Figures**

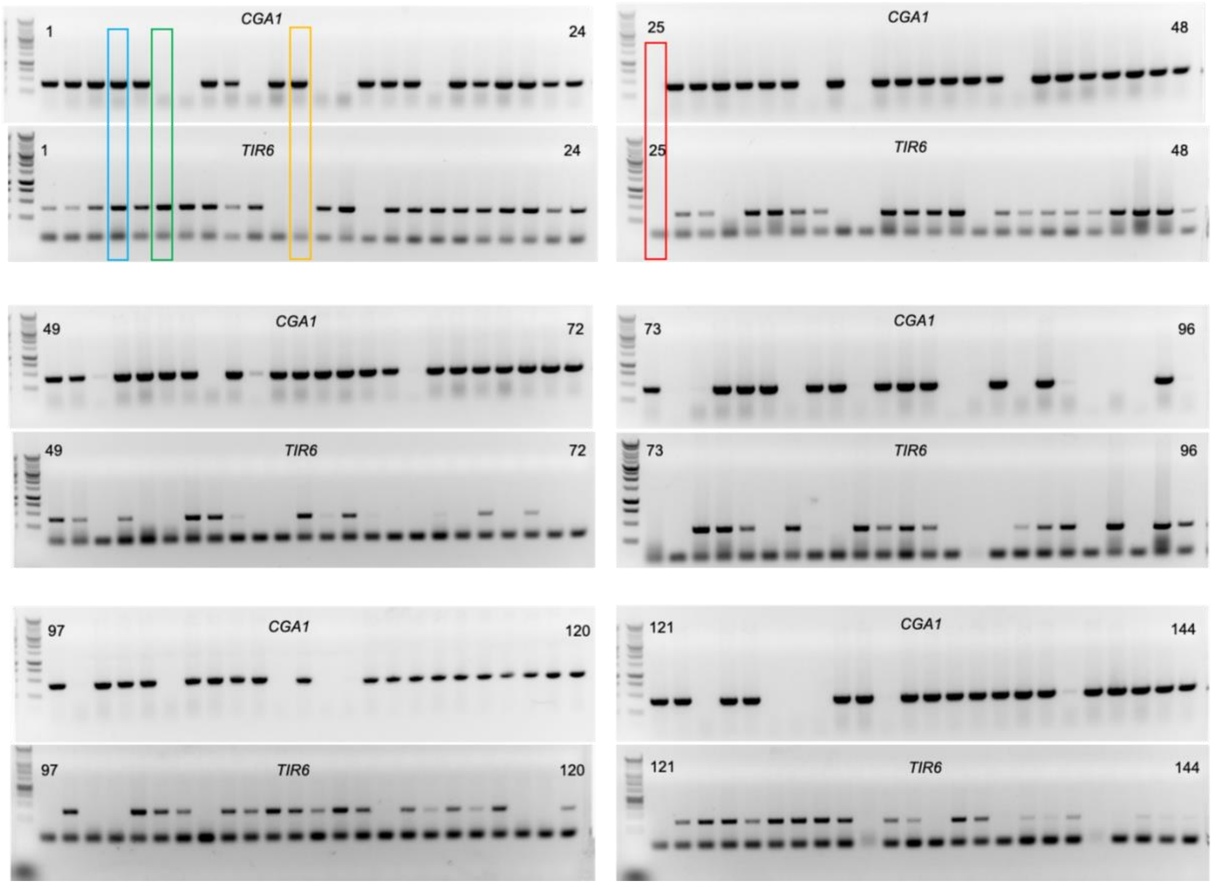


**Figure S1. PCR amplification of the *Cga1* gene and a combination of the *Cga1* gene and the transposon *TIR6* sequence from a B73 x 0634 maize $F_2$ progeny**. PCR reactions amplifying *Cga1* are displayed above the corresponding *TIR6* reactions, which involve a primer complementary to the *Mu* transposon terminal inverted repeat. Plants are in sequence, with the plant number for the beginning and end of each range displayed on the left and right side of each gel. Plants for which both PCR reactions were successful are heterozygous for a transposon insertion (plant #4 is shown as an example with a blue box). Plants for which the *Cga1* PCR is positive and the *TIR6* is negative are homozygous for wildtype *Cga1* (plant #12 is shown as an example with an orange box). Plants in which the *Cga1* PCR is negative and the *TIR6* PCR is positive have homozygous transposon insertions in *Cga1* (plant #6 is shown as an example with a green box). Plants in which both PCRs were negative indicate poor DNA quality, and were not used in further experiments.

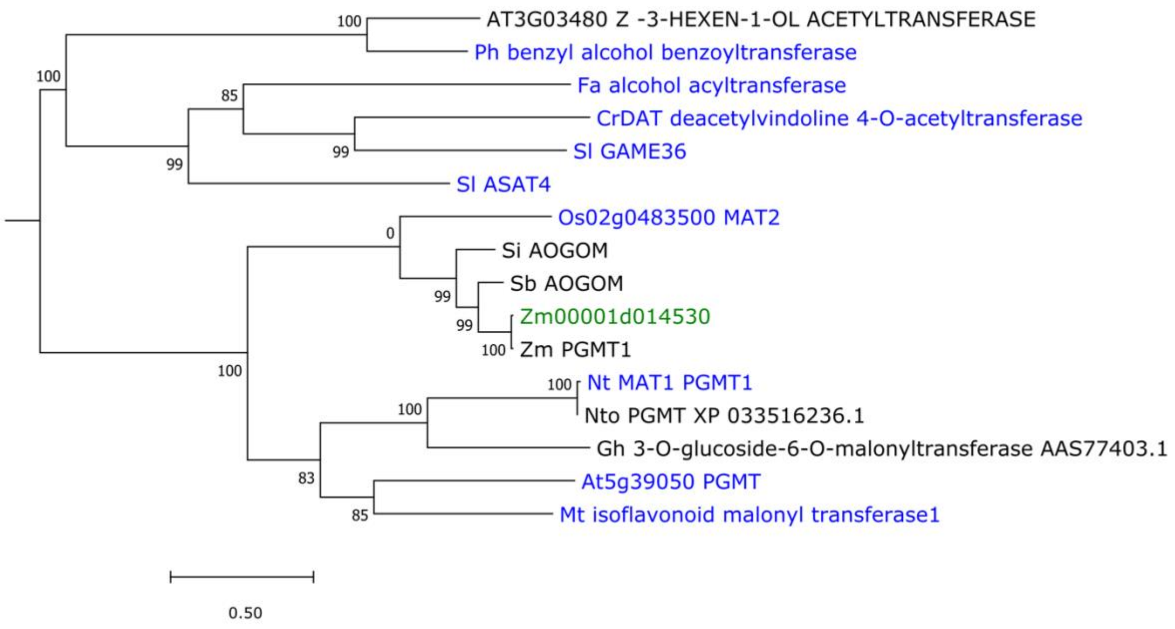


**Figure S2. Phylogenetic tree of acetyltransferases and malonyltransferases with different substrates.** Zm00001d014530 (PWZ24778.1, maize catechol glucoside acetyltransferase, CAG1) clusters with enzymes that acetylate or malonylate sugars connected to phenolic compounds. Green coloring indicates the acetyltransferase characterized in this study. Blue coloring indicates proteins whose function was determined experimentally, whereas black coloring indicates proteins whose function was attributed due to sequence homology. OsMAT2 (BAD21783.1)– a rice flavonoid malonyltransferase, that malonylates the 6”-hydroxyl group of the sugar (Kim *et al.*, 2009). Si AOGOM (XP_004969567.1)– *Setaria italica* anthocyanidin 5-O-glucoside-6''-O-malonyltransferase. Sb A0G0M (XP_002438902.1)-*Sorghum bicolor* anthocyanidin 5-O-glucoside-6''-O-malonyltransferase. Zm PGMT1 (AQK66481.1) – *Zea mays* phenolic glucoside malonyltransferase1. Nt MAT1 (NP_001312260.1)– *Nicotiana tabacum* malonyl transferase, that malonylates 2-naphthol-O-glucoside on position 6 (Taguchi *et al.*, 2010). Nto PGMT (XP_033516236.1)- *Nicotiana tomentosiformis* phenolic glucoside malonyltransferase. Gh 3-O-glucoside-6-O-malonyltransferase1 (AAS77403.1)– a *Glandularia x hybrida* enzyme. At5G39050 PGMT (NP_568561.4) – Arabidopsis phenolic glucoside malonyltransferase, which malonylates 2-naphthol-O-glucoside on position 6 (Taguchi *et al.*, 2010). Mt isoflavonoid malonyl transferase1 (ABY91220.1)– a *Medicago truncatula* enzyme that malonylates a sugar connected to an isoflavone (Yu *et al.*, 2008). At3G03480 (Z)-3-hexen-1-OL acetyltransferase (NP_186998.1)– an Arabidopsis enzyme predicted by homology to acetylate alcohols (D'Auria, 2006). Ph benzyl alcohol benzoyltransferase (Q6E593.1) – a petunia enzyme that can acetylate or add a benzoyl group to benzoyl alcohols (Boatright *et al.*, 2004). CrDAT deacetylvindoline 4-*O*-acetyltransferase (Q9ZTK5.1) – a *Catharanthus roseus* enzyme that catalyzes the last step in vindoline biosynthesis by acetylating a hydroxyl of deacetylvindoline (St-Pierre *et al.*, 1998). Sl GAME36 (XP_004246214.1) – a tomato enzyme that acetylates hydroxytomatine to form acetoxytomatine (Sonawane *et al.*, 2023). Fa alcohol acyltransferase (AAG13130.1) - a strawberry enzyme that acetylates short alcohols (C2-C7) (Olías *et al.*, 2002). Sl ASAT4 (NP_001266253.1)– a tomato acyltransferase, that acetylates sucrose in the biosynthesis of acylsugars (Fan *et al.*, 2016). Bootstrap values are based on 1000 replicates. Scale bar indicates substitutions per site.

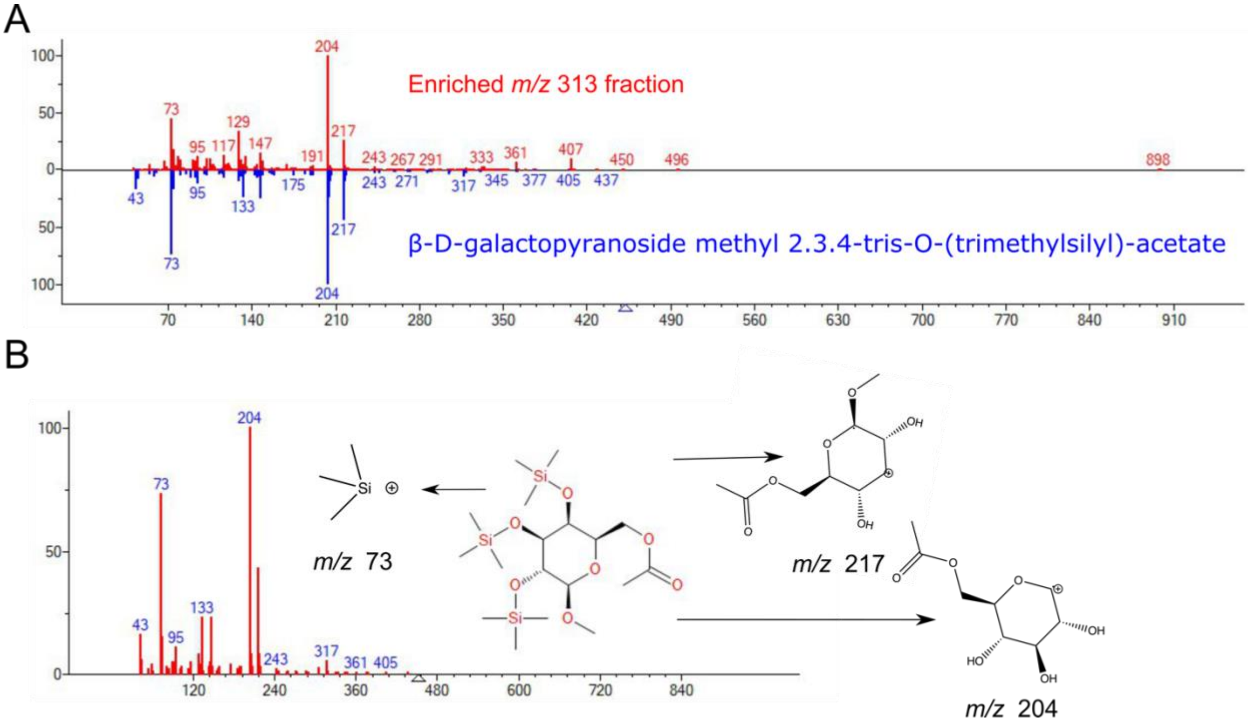


**Figure S3. Gas chromatography-mass spectrometry analysis of the *m/z* 313.09 compound fragmentation.** (A) Comparison of the fragmentation pattern of a fraction enriched in the *m/z* 313.09 compound (in red) and β-D-galactopyranoside methyl 2.3.4-tris-O-(trimethylsilyl)-acetate (in blue), as generated by the NIST spectral library. (B) the fragmentation and structure of β-D-galactopyranoside methyl 2.3.4-tris-O-(trimethylsilyl)-acetate. Three possible structures of the major fragments are presented.

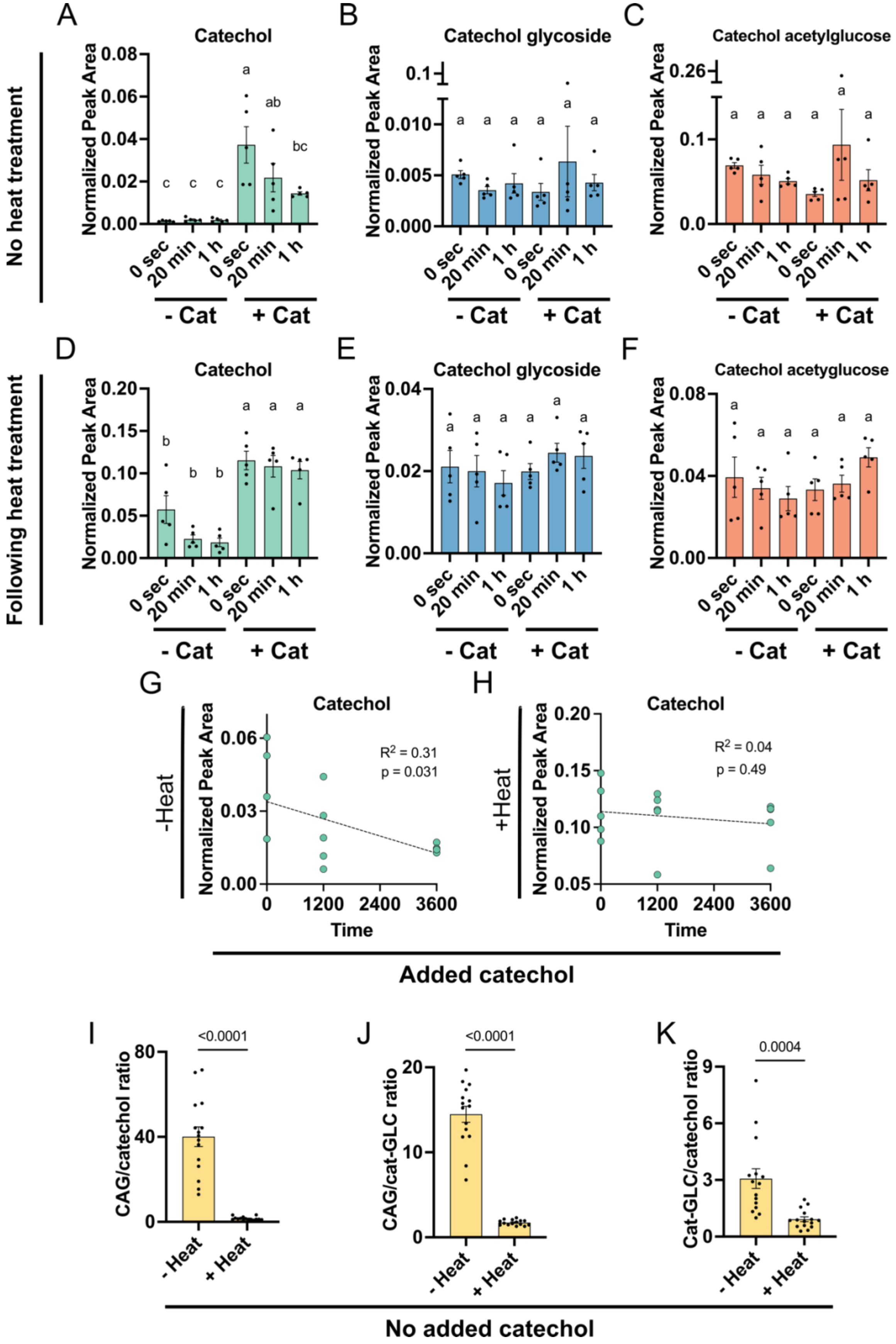

A
Catechol
B
Catechol glycoside
C
Catechol acetylglucose
No heat treatment
Normalized Peak Area
- Cat
+ Cat
0 sec
20 min
1 h
D
Catechol
E
Catechol glycoside
F
Catechol acetyglucose
Following heat treatment
G
Catechol
-Heat
R$^2$ = 0.31
p = 0.031
Time
H
Catechol
+Heat
R$^2$ = 0.04
p = 0.49
Added catechol
I
CAG/catechol ratio
<0.0001
J
CAG/cat-GLC ratio
<0.0001
K
Cat-GLC/catechol ratio
0.0004
- Heat
+ Heat
No added catechol

**Figure S4. Effects of incubation time and heat inactivation on the abundance of catechol metabolites, with and without catechol supplementation.** (A-C) Effects of incubation time on non-heat-treated samples. (A) Catechol. (B) Catechol glucoside. (C) catechol acetylglucoside (CAG). (D-F) Effects of incubation time on heat-treated samples. (D) Catechol. (E) Catechol glucoside. (F) CAG. (G-H) Correlation between incubation time and catechol abundance of catechol supplemented non-heat-treated (G) and heat treated (H) samples. (I-K) The ratio of different catechol derivatives in non-heat-treated and heat-treated samples. (I) ratio of CAG to catechol. (J) Ratio of CAG to catechol glucoside. (K) Ratio of catechol glucoside to catechol. (A-H) N=5. (I-K) N = 15. (A-F) Different letters indicate significance of $p < 0.05$, as determined by a Tukey's HSD test. (G,H) P values in scatter plots indicate significance of slopes' equality to 0, as determined by a regression *t*-test. (I-K) Values above bars indicate p values as determined by a two-tailed Student's *t*-test
