## Supplemental data S1 for "Catechol acetylglucose: A newly identified benzoxazinoid-regulated defensive metabolite in maize"

>Zm00001d014530_[PWZ24778.1]

MAVSPDQQQPPAGPGSPLPSPRLRVRVLDTTLVPPSPSPPETSLPLTFLDIFWLHSPPVERLFIYRLAPDADVDAIISRLRDSLHQAVRAFYPLAGRLRLTPGTSDRYELHYRPGDAVTFTVAECDDDDADAHFDALATDEPREVAKIATLVPPLPERGGLFAVQATLLPARRGLSIGVTVHHAACDGSGSTHFLHTWAVACRAGAEAPPPPPVIDRSLIADPRRMYDVFIQGAPSTEEYEFVKMSADQLFATFALSKDELKHIKDAVAEEAARRGVAPPRCSSLVATFGFVWSCYQQAKEASSGAGEFPMTCMAFPVDHRSRMEPPLPDKYLGNCVGPAFALAPTGELAQAGAGGFFSACAAVASSIDEAVRDIETSNVEVWFDRIRDLIPMGFLTVAGSPRFRVYDMDFGFGRPAKVDVVSVARTGAVAVAESRDGNGGIEVGVSLQPAAMKRYRKCFADAIAWLHQRT

>Zm_PGMT1_[AQK66481.1]

Mddatyrpedqcdgpiathcltqarssksdvdtigrcaamavspdqqqppagvgsplpsprlrvrvrdttlvppspsppetslpltfldifwlhsppverlfiyrlapdadvdaiisrlrdslhqavrafyplagrlrltpgtsdryelhyrpgdavtftvaecddddadahfdalatdeprevakiatlvpplpergglfavqatllparrglsigvtvhhaacdgsgsthflhtwavacragaeappppppvidrsliadprrmydvfiqgapsteeyefvkmsadqlfatfalskdelkrikdavaeeaarrgvapprcsslvatfgfvwscyqqakeassgagefpmtcmafpvdhrsrmepplpdkylgncvgpafalaptgelaqagaggffsacaavassideavrdietsnvevwfdrirdlipmgfltvagsprfrvydmdfgfgrpakvdvvsvartgavavaeswdgnggievgvslqpaamkryrkcfadaiawlhqrt

>Sb_AOGOM_[XP_002438902.1]

Mamapdqqqppagagagassttgstsprlrvhvhdttlvppspsppetslpltffdvfwlqshpverlflyrlahdadveaiisnlrsslhkalaafyplagrvrltpgtsdryelhyrpgdavtftvaecdddvdgdahfdalatdeprevakiaalvptlprggrllavratllparrglaigvtlhhaacdgsgsthflhtwaatcrgggaesppppvidrtlladprrlydafvqtapsseeyefvkmsadqlfatftlskddlkrvkdavadeaarrgvapprcsslvatfglvwscyqrgkegsgggagegsmacmafpvdhrsrmkpplpekylgncvgpafalaptgelaaagagglfsacaavasaideavrdigtssmdawmdrirevlpmglltvagsprfrvydldfgfgrpakvdivsvartgavavaesrsgdggievgvslqpaameryrkcfadatlwlhqkt

>Si_AOGOM_[XP_004969567.1]

mavapdqkqrpagsssasphlrvhdtilvapspsppetslpltffdiiwlnsppverllfyrlapdadvatiisnlkdslhkafrvfyplagrlrltpgtsdryelyyspgdavtftvaecddgdadidglaagdprevakigtlvpplaeggglfalqatllsarrglaigvtvhhaacdgsnsthflhtwaaacsgteappppvidrtlladprglynvfyqeapstdemefakmsadqlfatftlskddlqrikevvadeaarrgvapprcsslvatfgfvwscyqrakescgsgegpmtcilfpvnhrsrmkpplperylgncvgpafgmapkselavagvgglftacaavasaideavrdigtssmdawldrikeasangilsvagsprfrvyeldfgfgrplkvdivsvartgavavaesrsciggmevgvslqpagmdryrkcftdgiawlhqrs

>At_At5g39050_PGMT_[NP_568561.4]

MVNEEMESSLKVIDVARVTPSNSDSSESLTLPLTFFDLLWYKLHAVERVIFYKLTDASRPFFDSVIVPNLKTSLSSSLSHYLPLAGKLVWEPLDPKPKIVYTPNDAVSFTVAESNADFSRLTGKEPFPTTELYPLVPELHVSDDSASAVSFQVTLFPNQGFCISVNAHHAVLDGKTTTNFLKSWARTCKNQDSFLPQDLIPVYDRTVIKDPMDLDTKILNAWHRVAKVFTGGKEPENPKSLKLLWSPEIGPDVFRYTLNLTREDIQKLRERLKKESSSSSVSSSPKELRLSTFVIVYSYALTCLIKARGGDPSRPVGYGFAVDCRSLMVPPVPSSYFGNCVSACFKMSLTAETFMSEEGFLAAARMVSDSVEALDENVALKIPEILEGFTTLSPGTQVLSVAGSTRFGVYGLDFGWGRPEKVVVVSIDQGEAISFAESRDGSGGVELGFSLKKHEMDVLVDLLHKGLEN

>Nt_MAT1_PGMT1_[NP_001312260.1]

MASVIEQCQVVPSPGSATELTLPLTYFDHVWLAFHRMRRILFYKLPISRPDFVQTIIPTLKDSLSLTLKYYLPLAGNVACPQDWSGYPELRYVTGNSVSVIFSESDMDFNYLIGYHPRNTKDFYHFVPQLAEPKDAPGVQLAPVLAIQVTLFPNHGISIGFTNHHVAGDGATIVKFVRAWALLNKFGGDEQFLANEFIPFYDRSVIKDPNGVGMSIWNEMKKYKHMMKMSDVVTPPDKVRGTFIITRHDIGKLKNLVLTRRPKLTHVTSFTVTCAYVWTCIIKSEAATGEEIDENGMEFFGCAADCRAQFNPPLPPSYFGNALVGYVARTRQVDLAGKEGFTIAVELIGEAIRKRMKDEEWILSGSWFKEYDKVDAKRSLSVAGSPKLDLYAADFGWGRPEKLEFVSIDNDDGISMSLSKSKDSDGDLEIGLSLSKTRMNAFAAMFTHGISFL

>Nto_PGMT_[XP_033516236.1]

masvieqcqvvpspgsateltlpltyfdhvwlafhrmrrilfyklpisrpdfvqtiiptlkdslsltlkyylplagnvacpqdwsgypelryvtgnsvsvifsesdmdfnyligyhprntknfyhfipqlaeskdapgvqlapvlaiqvtlfpnhgisigftnhhvagdgativkfvrawallnkfggdeqflanefipfydrsvikdpngvgmsiwdemkkykhmmkmsdvvtppdkvrgtfiitrhdigklknlvltrrpnlthvtsftvtcayvwtclikseaatgeeidengmeffgcaadcraqfnpplppsyfgnalvgyvartrqvdlagkegftiaveligeairkrmkdeewilsgswfkeydkvdakrslsvagspkldlyaadfgwgrpeklefvsidnddgismslskskdsdgdleiglslsktrmnafaamfthgisfl

>Sl_Solyc08g075210.1.1_GAME36_[XP_004246214.1]

MTASSFVSMAEKIIKPHSPTPFSVKRYNLCLMDEIMVPVYMPIVAFYPNPSKTPEQVSNILEDSLSKVLSSYYPFAGTLGSDNATFVDCNDRGAKSIQVRYDCPMSEIVNLPDTGPEYLPFAKGTPWSSTPEEQSLLVVQLSHFNCGGLGISARLSHKIADGCTLANFISDWASVARDDNANIPSPQLIGSSIFPPFTEMRIHTDTNVDYEFYNLPVCKKRYLFSNAKLEMLKTQVESETGVQNPTRIEVLSALIYKCAVTANSSSFRPSSLSLPVNLRPILNPPLETRTVGNIISFIKVETTSEDEMTIGRVVREIRKGKDELKQEGGVKKEKLVSLWSEWIHSIDLYRSSSVCNYPLNNLDFGWGKPNRVAIPVFGVANTCMFMDNLSGDGIEVIIALPEKDATQFENSKELLHFASPVTNL

>AT3G03480_(Z)-3-HEXEN-1-OL_ACETYLTRANSFERASE_[NP_186998.1]

MDHQVSLPQSTTTGLSFKVHRQQRELVTPAKPTPRELKPLSDIDDQQGLRFQIPVIFFYRPNLSSDLDPVQVIKKALADALVYYYPFAGRLRELSNRKLAVDCTGEGVLFIEAEADVALAELEEADALLPPFPFLEELLFDVEGSSDVLNTPLLLVQVTRLKCCGFIFALRFNHTMTDGAGLSLFLKSLCELACGLHAPSVPPVWNRHLLTVSASEARVTHTHREYDDQVGIDVVATGHPLVSRSFFFRAEEISAIRKLLPPDLHNTSFEALSSFLWRCRTIALNPDPNTEMRLTCIINSRSKLRNPPLEPGYYGNVFVIPAAIATARDLIEKPLEFALRLIQETKSSVTEDYVRSVTALMATRGRPMFVASGNYIISDLRHFDLGKIDFGPWGKPVYGGTAKAGIALFPGVSFYVPFKNKKGETGTVVAISLPVRAMETFVAELNGVLNVSKG

>CrDAT_deacetylvindoline_4-O-acetyltransferase_[Q9ZTK5.1]

mesgkisvetetlsktlikpssptpqslsrynlsyndqniyqtcvsvgffyenpdgieistireqlqnslsktlvsyypfagkvvkndyihcnddgiefvevrircrmndilkyelrsyardlvlpkrvtvgsedttaivqlshfdcgglavafgishkvadggtiasfmkdwaasacylssshhvptpllvsdsifprqdniiceqfptskncvektfifppeaieklkskavefgiekptrvevltaflsrcatvagksaaknnncgqslpfpvlqainlrpilelpqnsvgnlvsiyfsrtikendylnekeytklvinelrkekqkiknlsrekltyvaqmeefvkslkefdisnfldidaylsdswcrfpfydvdfgwgkpiwvclfqpyikncvvmmdypfgddygieaivsfeqekmsafekneqllqfvsn

>Fa_alcohol_acyltransferase_[AAG13130.1]

mekievsinskhtikpstsstplqpykltlldqltppayvpivffypitdhdfnlpqtladlrqalsetltlyyplsgrvknnlyiddfeegvpylearvncdmtdflrlrkieclnefvpikpfsmeaisderypllgvqvnvfdsgiaigvsvshklidggtadcflkswgavfrgcreniihpslseaallfpprddlpekyvdqmealwfagkkvatrrfvfgvkaissiqdeaksesvpkpsrvhavtgflwkhliaasraltsgttstrlsiaaqavnlrtrmnmetvldnatgnlfwwaqailelshttpeisdlklcdlvnllngsvkqcngdyfetfkgkegygrmceyldfqrtmssmepapdiylfsswtnffnpldfgwgrtswigvagkiesasckfiilvptqcgsgieawvnleeekmamleqdphflalaspktli

>Mt_isoflavonoid_malonyl_transferase1_[ABY91220.1]

masnnnsnikvhdhfkvvppsstkttsipltffdifwlrfhpvervffytlpnsqshpsfffqtivpnlksslsltlqhflplagnivwpsdsskpiiqfdpnddgvsliiaesdsdfnhvvenspheaslsrsfiphlessdsfasimslqitlfpnsgfsigisthhavldgksstmfvkawaylckkaierdesptllsefepsfnrevikdpngnnvmdlvstlfpsekgndrslkifpfepqledsvratfklkhedldkikqrvlstweifdtkeskpqtlssfvitcayslvcvakaihgahndkekfsfvfsvdcrarleptipnnylgncvwayfidtqpldfikedgvflvaksiyekikminekgflegeindmfnkiislssegfefmgvagshrfgvyeidfgwgrpekveivsidrgvtiglaeskdskggievglalnkpvmdifstlfleglsyne

>Ph_benzyl_alcohol_benzoyltransferase_[Q6E593.1]

mdskqsselvftvrrqepeliapakptpretkflsdiddqeglrfqipvinfyrkdssmggkdpvevikkaiaetlvfyypfagrlregndrklmvdctgegvmfveanadvtleefgdelqppfpcleellydvpgsagvlhcpllliqvtrlrcggfifalrlnhtmsdapglvqfmtavgemargatapstlpvwcrellnarnppqvtcthheyeevpdtkgtliplddmvhrsfffgptevsalrrfvpphlhncstfevltaalwrcrtisikpdpeeevrvlcivnarsrfnpqlpsgyygnafafpvavttaeklcknplgyalelvkktksdvteeymksvadlmvikgrphftvvrtylvsdvtragfgevdfgwgkavyggpakggvgaipgvasfyipfrnkkgengivvpiclpgfamekfvkeldsmlkgdaqldnkkyafitpal

>Sl_acylsugar_acetyltransferase_AT2_[NP_001266253.1]

mncyieiqsrkmvkpsaptpdnlrrlklslfdqmdigayvpivfnylpnstssydhddklekslsetltkfypfagrfrkgidpfsidcndegieyvrtkvnaddlaqylrgqahndiesslidllpvmhrlpssplfgvqvnvfnnggvtigiqilhmvsdaftlvkfvnewahttltgtmpldnpgfgqlpwlfparalpfplpdfntttapnyknvtkrflfdalaienlrntikandmmmkqpsrvvvvmsliwkvlthissaknngnsrdsslvfvvnlrgklsctapslehvvgncvipatankegdearrkddelndfvklvrntirdtceaigkaesvddisslafnnltkciekilhgdemdfyscsswcgfpwyeadfgwgkpfwvssvsfghhgvtnlmdtkdgdgiqvticlkendmieferdphilsstsklafhslg

>Gh_3-O-glucoside-6-O-malonyltransferase_[AAS77403.1]

Matipkpatvlercrvapapppngsdaaehpqtlplvfsdmvwlhfhptqrllfykfpcskdhfiehivpnfkkplsqalkhflpfagnliyeidsgnipelqyrpgdsvpvtvaesneasdfdylsgdrprdagefyaffpdfppdkiesgwkkipllavqitlfpdtgicigfnnnhivgdassvvgfikawssicrhggdeefssathlhpfydrsvikdptgllnnywnqmkhvkiespplnfptnklratyilqksdiqnlrnlvqaqkqdlvhlssftittayvwtclakssaaageevaadeteyfvfavdarqrinppapatyfgncvvpafaesthselkgkngfltavglvgdviskkannkdeilrgadewlaklggmigkrlfgvagspkfdlydadfgwgnpnkyesvgidvdgsmslcksrefeggleiglslpkkkmdafgdvfcgglkiteqts

>Os02g0483500_MAT2_[BAD21783.1]

MAPATQMAAPPPRARGGSFRVLRTARVAPSSPDGVPMLVERAVPLTFLDAIWLPTPPVDRVFFYRLGADDDGVDAVLSRLADSLSRALHVVYPLAGRLRLTPGKTNRYELFYQPGDAVAFTFAEHDDGVGVDELAADDPREVAKIAPLVPELPDGGAVLAVQATVLPPARRGLALGVTVHHAACDGSSSTHFLHTWAAACAGAAVLPKPPVIDRTFIREREDLYDYMVSRTEESDKFRSPDVADSKLLATFTLSGEILQSIKDRVAGVAARRGKSPPPRCTSVVATFAIVWQCHIRAALGDVEADNKHHGRAHFIFPTDHRARMEPRVPDKYLGNCVGPCFASAPKEEIAAADAEDGLYTTCAAIAAAVDEGTRYDPDYWKRCMEHVGGMSASDGPPLAVAGSPRFRVYDVDFGFGRPAKVDVVSVAKTGAISVAEGRRGGIEVGVGLPPERMERFRRCFADAVAWLSSPSRPVTRDMDRSAPGHSPA
